# Separate timescales for spatial and anatomical information processing of body stimuli

**DOI:** 10.1101/2024.10.07.616993

**Authors:** Baptiste M. Waltzing, Siobhan McAteer, Marcos Moreno-Verdú, Elise E. Van Caenegem, Yue Du, Robert M. Hardwick

**Author notes:** Correspondence should be addressed to Baptiste Waltzing.

## Abstract

Observing different body stimuli can influence the speed and accuracy of our responses. Prior work indicates this effect is influenced by factors such as spatial congruence and perspective. We hypothesized that the influence of these factors would vary depending on the amount of time that participants had to process visual stimuli. Experiment 1 was a reaction time task (n=29) with stimuli varying in spatial congruence (congruent, incongruent, neutral), perspective (first- or third-person) and stimulus type (body or control). Experiment 2 (n=50) used the same stimuli in a “Forced Response” paradigm, which controlled the time participants had to prepare a response. This allowed us to assess responses as a function of preparation time. Experiment 1 showed effects of spatial congruence, with longer reaction times and more errors for spatially incongruent stimuli. This effect was greater for body stimuli. Experiment 2 showed that spatial information was processed faster than anatomical information, inducing incorrect responses at short preparation times for spatially incongruent body stimuli. There was little-to-no corresponding effect for control stimuli. Both experiments also showed weak-to-no effects of perspective, which appear to have been driven by spatial congruence. Our results indicate that spatial information is processed faster than anatomical information during observation of body stimuli. These data are consistent with the dual visual streams hypothesis, whereby spatial information would be processed rapidly via the dorsal stream, while anatomical processing would occur later via the ventral stream. These data also indicate differences in processing between body and control stimuli.

**Public significance statements:** This study provides novel insight into the time-course of information processing, showing that spatial information is processed faster than anatomical information for body stimuli. The results also challenge the established view that visual perspective is critical to process body stimuli, demonstrating that this may instead result from lower-level effects of spatial congruence.

## Introduction

Action observation (i.e. observing a movement being performed) plays a central role in our everyday lives. We observe the movements of others in many contexts, including to imitate them in order to learn new movements (Gatti et al., 2013), to respond with a complementary action such as shaking hands (Curioni et al., 2020; Faber et al., 2016), or without any specific intention. In recent years, action observation has also been used in the medical field, notably for the rehabilitation of movement-related disorders (Caligiore et al., 2017; López et al., 2019; Machado et al., 2019; Nicholson et al., 2019). Despite this widespread interest, there are still fundamental questions regarding how stimuli presenting actions are processed; addressing these questions is a key challenge in optimizing the use of action observation in order to use it as effectively as possible in applied situations.

In non-human primates, the effects of action observation are mediated by mirror neurons. These neurons, first discovered in the F5 region of the brains of macaques, are activated both during the observation of an action, and during the execution of the same action (di Pellegrino et al., 1992; Rizzolatti et al., 1996). Neurons with similar properties have been identified in premotor and parietal areas, and are thought to form a “mirror neuron system” that could mediate the effects of action observation (Blakemore & Frith, 2005). In humans, action observation leads to the recruitment of functionally similar premotor and parietal areas that are considered to form an “action observation system” (Fadiga et al., 1995; Hardwick et al., 2018). These discoveries provide a potential neural basis for behavioural effects whereby observing the actions of another person can influence our own movements.

Research has demonstrated that observing actions performed by other people can prime us to perform similar movements (Edwards et al., 2003; Hardwick & Edwards, 2011, 2012; Press et al., 2011). In particular, previous work indicates that reaction times are reduced following the observation of an action compatible with the required response (Brass et al., 2001, Edwards et al. , 2003). Subsequent studies have investigated factors that influence such priming effects (for a review see (Kemmerer, 2021)). In this study, we focused on three of them in particular. The first is the nature of the stimulus, which may be body-related, or control. Studies have shown that the observation of body-related stimuli (e.g. a hand (Gowen et al., 2010)) can induce a priming effect if the requested movement is congruent with the observed movement. There is mixed evidence that this priming effect is smaller or absent during the observation of control stimuli (e.g. geometric shapes (Biermann-Ruben et al., 2008)) and is therefore argued that there would be a particular processing for body stimuli (for a review see (Gowen & Poliakoff, 2012)). This greater priming effect could be explained by the pre-activation of motor areas when the action is observed (Jeannerod, 2001). Many different control stimuli have been used in past studies, e.g. squares, pens, dots, crosses, etc; for review see (Gowen & Poliakoff, 2012). While numerous possible control stimuli were possible, each with respective strengths and limitations, our choice in the present study was conditioned upon certain requirements. A primary point was that we required control stimuli that were asymmetrical, so that they could undergo vertical and horizontal symmetries equivalent to a change in laterality or perspective of the observed hand. We also required the stimuli to be easy to associate with the required response, so that extensive training would not be necessary. Some of the control stimuli previously used were superimposed on hand images (e.g. X superposed on finger)(Brass et al., 2001). As we wanted stimuli that didn’t include a body element, this choice was not possible. We therefore chose to use the letters of the keyboard buttons that participants would press to act as the control stimuli in the present task, as they fulfilled the criterion noted above. A second factor is perspective. It has been shown that observing an action from a first-person perspective (consistent with observing our own movements through our own eyes) had a greater priming effect and was therefore associated with faster reaction times than actions observed from a third-person perspective (observing an action from the vantage point of another person) (Mibu et al., 2020). This effect could be explained by the fact that some studies have shown greater activation of areas of the action observation system when observing actions from a first-person perspective than from a third-person perspective (Angelini et al., 2021; Ge et al., 2018; Oosterhof et al., 2012). The third factor we were interested in was the spatial congruence of the observed action. Studies have shown that observing an action that is spatially congruent with the required action (e.g. a right hand in first-person perspective for a response with the right hand) (fig 1A) has a greater priming effect than observing an action that is spatially incongruent (e.g. a left hand in first-person perspective for a response with the right hand) (Avikainen et al., 2003; Bekkering et al., 2000; Chiavarino et al., 2007; Wohlschläger & Bekkering, 2002). This could be explained by greater activation of the contralateral hemisphere of the hand observed in the first-person perspective but greater activation of the ipsilateral hemisphere in the third-person perspective (Shmuelof & Zohary, 2008).

**Figure 1.**
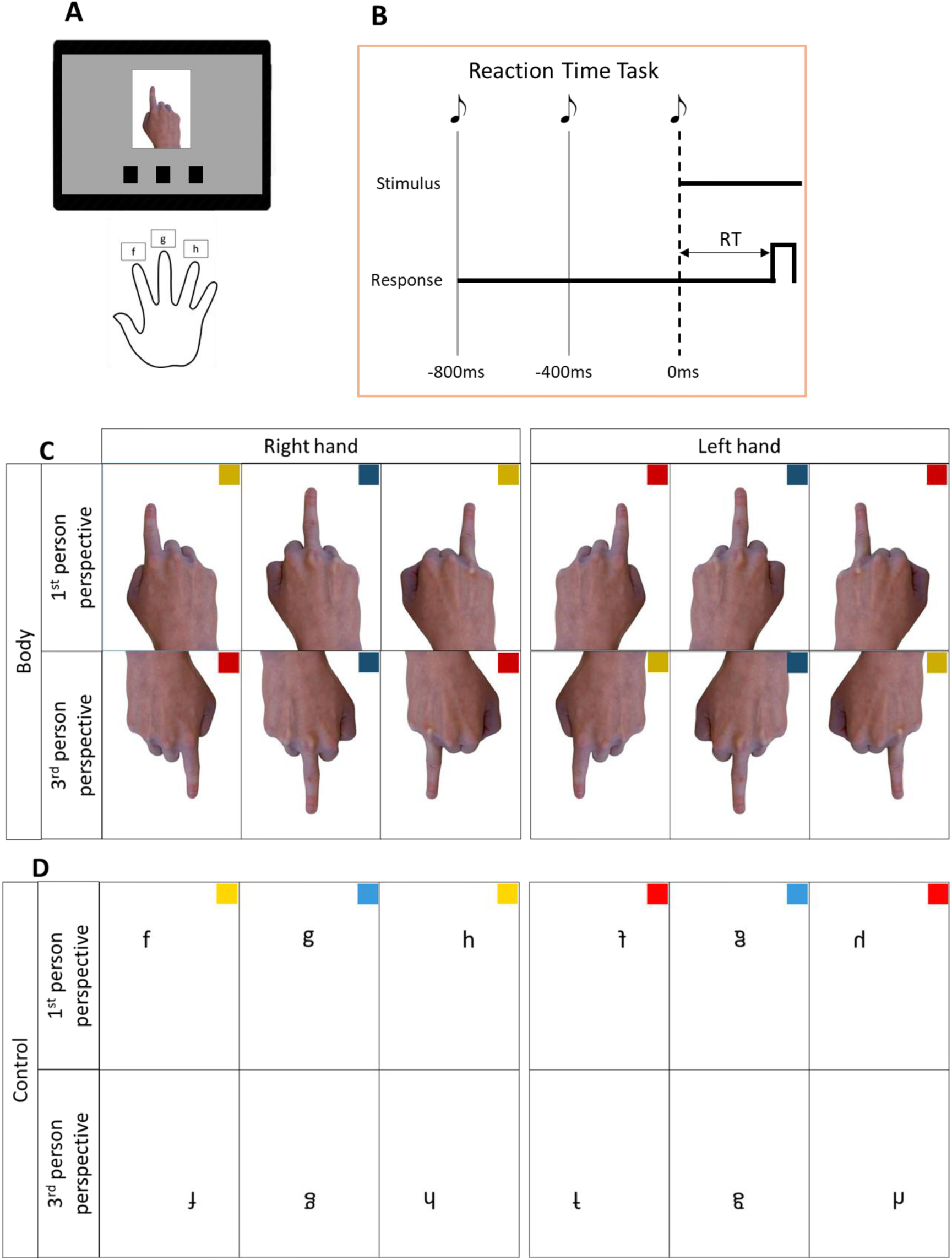
A. General experimental task set up. Participants positioned their right hand with the index, middle and ring fingers resting on the f, g and h keys respectively. The stimulus appeared in the centre of the screen, indicating which key should be pressed. Underneath were 3 black squares, each corresponding to a key and providing visual feedback. The square of the key pressed turned green if the response was correct and red if the response was incorrect. B. Reaction time paradigm. In each trial the participant heard a sequence of three equally spaced tones, and a stimulus always appeared at the same time as the third tone. Participants were instructed to respond as quickly and accurately as possible once the stimulus appeared. C. Body stimuli. The different body stimuli are presented, which can show a right or left hand, first- or third-person perspective, with index, middle or ring finger raised. The coloured squares are not part of the stimuli, but indicate the spatial congruence: yellow for spatially congruent stimuli, red for spatially incongruent stimuli and blue for spatially neutral stimuli. A darker shade is used for body stimuli and a lighter shade for control stimuli. This colour code is reused in the results plots. D. Control stimuli. These stimuli were presented in the same position as the body stimuli, whose low-level characteristics they mimic.

We therefore examined how these three factors influenced action observation. Experiment 1 used a classical reaction time task to study possible interactions between the factors of stimulus type, perspective and spatial congruence. We predicted that although perspective may have an effect, the most important point would be spatial congruence, with shorter reaction times for spatially congruent stimuli than for spatially incongruent stimuli. This hypothesis is inspired by the work of Simon et al. (Simon & Wolf, 1963) who showed that the spatial position of the observed stimuli played a key role in their processing, with shorter reaction times when they were spatially congruent. While traditional reaction time-based studies provide general information about stimulus processing, they are limited in their ability to examine the precise time-course of information processing. To address these limitations, Experiment 2 used a forced response paradigm (Haith et al., 2016; Hardwick et al., 2019; Schouten & Bekker, 1967; Vleugels et al., 2020). This procedure limits the processing time available to participants, providing a measurement of accuracy as a function of the response preparation time available, allowing us to obtain a novel insight into how the stimulus is processed. We expect to observe differences in processing between body and control stimuli. We therefore hypothesize that, due to the activation of the action observation system, which includes motor-related areas specifically following the observation of body stimuli (Hardwick et al., 2018) body-related stimuli would be more susceptible to changes in spatial congruence. Similar to our prediction for Experiment 1, we also hypothesised that spatial congruence may interact with perspective during this task.

## Experiment 1

### Experiment 1 Methods

In this first experiment, we investigated how action observation could be influenced by factors including the nature of the stimulus, perspective, and the spatial congruence of the stimulus with the participant’s responses. To do this, we performed a reaction time task to examine variations in both reaction time and errors. All procedures were reported in accordance to the ‘Action Observation’ part of the “Guidelines for Reporting Action Simulation Studies” checklist (Moreno-Verdú et al., 2024). This study was not preregistered.

### Participants

A group of 30 healthy young participants were recruited via the Prolific online platform (https://www.prolific.com/). A single participant was excluded for an extremely high number of unanswered or incorrect trials and very short reaction times (see section 1.1 of appendices for more details). This left 29 participants aged between 18 and 34 years (mean = 25.3 years, SD = 4.2 years). All participants were right handed, 10 were women, 19 were men. The study by Catmur and Heyes, 2011 (Catmur & Heyes, 2011), suggests a moderate effect size (dz = 0.71)(Cohen, 1977) for the influence of spatial congruence on reaction time. A sample size calculation with Gpower 3.1 for a paired two tails t-test with alpha = 0.05, power = 0.8 and a moderate effect size (dz = 0.71) suggests that a sample size of more than 18 participants would be sufficient. To ensure sufficient power in case of additional variability, the number of participants was increased to 30. They were all naive about the purpose of the experiment. All participants gave their consent before the start of the experiment and were financially compensated (3£) for their time. The study was approved by the IPSY ethics committee of UCLouvain.

### General Task Procedures

The task was created in PsychoPy2 version 2023.2.2 (Peirce et al., 2019) and run online via PsychoPy2’s associated webpage Pavlovia (https://pavlovia.org/) (Bridges et al., 2020).

This reaction time task consisted of three different conditions: the familiarisation condition, the body stimuli condition, and the control stimuli condition. The task was the same for all three conditions, but each condition was characterised by different stimuli. Participants sat in front of a computer keyboard with the index, middle, and ring fingers of their right hand positioned on the f, g, and h keys of the keyboard, respectively (fig 1A). For each trial, the participants heard three short tones (50ms duration) spaced 400ms apart (fig 1B). To make it easier to distinguish between the different tones, the first two tones had a lower pitch than the final tone (i.e. notes “La” and “Si”, respectively). A visual stimulus was presented at the same time as the third tone. Depending on the stimulus that appeared, the participants had to respond by pressing on the keyboard with their index, middle, or ring finger. Participants were instructed to answer as quickly and as accurately as possible once the stimulus appeared on the screen. They were given a maximum of 1s to respond, or the trial was considered to be ’time out’. Once they had answered, visual feedback was displayed on the screen to indicate whether or not the answer was correct. The feedback consisted of three small black squares at the bottom of the screen, each corresponding to one of the three keyboard buttons. The box corresponding to the button that the participant responded with changed to provide feedback, turning green if the response was correct, or red if it was incorrect. The trial had a fixed duration of 1 second regardless of the participant’s reaction time, and was followed by a 200ms interval in which no stimulus was shown on screen. In all conditions the order of trials was pseudo-randomised such that the same response was presented on a maximum of two consecutive trials. There was a break between each block (this could be as long as the participant desired, with a minimum of at least 10s), during which the instructions were displayed on the screen. Each block began with a short check in which the participant had to press the three keys, in order to prompt the participants to ensure their fingers were placed on the correct keys.

### Familiarisation condition

The familiarization block was used to help participants gain some practice in completing the task. For this condition, one of three different stimuli could be displayed on the screen: an arrow pointing to the left, upwards, or to the right. For these stimuli, participants had to respond with their index, middle and ring fingers respectively. This block consisted of 20 trials for each stimulus. Data from this condition was not included in later analyses.

### Body stimuli condition

For this condition, the stimuli displayed were images of a hand with one finger raised (fig 1C). Participants were instructed to respond using the finger that appeared raised on the displayed stimulus (i.e. if the stimulus presented an index finger, the participant had to respond with their own index finger by pressing on the f key). The hand could be presented in first- or third-person perspective, be a right or left hand, and have the index, middle, or ring finger raised, for a total of 12 different stimuli (2 perspectives * 2 hands * 3 fingers). The same three images of the right hand in first-person perspective, one for each finger, were used for all stimuli, with mirroring and/or rotation to create left hand/third-person perspective stimuli. The position of the stimulus relative to the participant’s own hand, and thus the spatial congruence of the stimulus relative to the required response, was therefore influenced by the perspective and the hand observed. For example, when the stimulus presented the index finger of a right hand in the first-person perspective, the position of the finger was spatially congruent with that of the participant’s own index finger; by contrast, if the stimulus presented the index finger on a *left* hand in first-person perspective, the position of the finger was spatially incongruent with the response required by the participant. While these manipulations affected the positions of the index and ring fingers, being centred, the stimuli with the middle finger were always spatially congruent regardless of the perspective and the hand observed. For this condition, there were two blocks of 120 trials, with each stimulus being presented 10 times per block. The order of the trials was pseudorandomised to limit the number of consecutive responses with the same finger to two. Participants therefore completed a total of 20 trials with each stimulus, and a total of 240 trials in this condition.

### Control stimuli condition

Previous studies have suggested differences in processing for body compared to control stimuli (Gowen & Poliakoff, 2012). Therefore, the aim of this condition was to have control stimuli that mimicked the low-level characteristics of body stimuli, to see whether the effects observed were specific to action observation or not. The stimuli were the lowercase letters “f”, “g” or “h” (fig 1D) in Calibri font. Participants were asked to respond to these stimuli with their index, middle and ring fingers respectively. To obtain the 12 different stimuli equivalent to the body stimuli, the letter stimuli were first placed at the respective locations of the index, middle and ring fingers of the right hand stimulus in first-person perspective. The same rotation/mirroring manipulations as applied to the body stimuli were then applied to these control stimuli. This gave us 12 different stimuli (2 * rotation (equivalent to “perspective” in the body stimuli condition), 2 * vertical symmetry (equivalent to “hand” in the body stimuli condition), 3 * letters (equivalent to “finger” in the body stimuli condition). Like the body stimuli, the positions of the letters f and h could be spatially congruent (i.e. equivalent right hand, first-person perspective and left hand, third-person perspective) or incongruent (i.e. equivalent left hand, first-person perspective and right hand, third-person perspective). The central position of the letter g meant that it was always spatially congruent, regardless of the orientation of the stimulus. As in the body stimuli condition, participants completed two blocks, each block comprising 120 trials with 10 trials for each stimulus, for a total of 240 trials with the same pseudorandomisation.

### Protocol

Participants always began with the familiarisation condition, after which the order of conditions was counterbalanced across participants; half of the participants completed the body stimuli condition first, followed by the control stimuli condition, and vice-versa. The median completion time was 24 minutes (including breaks between blocks).

### Data analysis

Analyses were performed in R version 4.3.2 (R Core Team 2024).

We measured two variables of interest: reaction time (defined as the time between the appearance of the stimulus and the response given by the participants) and error rate (percentage of incorrect responses out of the total number of responses).

### Exclusion criteria

Of the 13920 total trials conducted, 370 trials (representing 2.66% of the total) were excluded from the analyses for having no response.

### Reaction time

As we performed a separate analysis for the error rate, trials with incorrect responses were not included in the reaction time analyses (n=1421), leaving a total of n=12129 trials for analysis.

### Global Analysis

Data were first analysed using a “Global” model that considered the data in its most granular form. A complex random intercept model (the choice of model was made following the guidelines in (Scandola & Tidoni, 2024)) examined effects and possible interactions between several factors, labelled here in relation to the body stimuli. This included the factors of “Stimulus Type” (2 levels; body or control), “Perspective” (2 levels; first-person or third-person), “Hand” (2 levels; left or right), and “Finger” (3 levels; index, middle, or ring). The model included between-participant variability as a random effect (i.e. random intercept) and used the formula: reaction time ∼ stimulus type * perspective * hand * finger + (1|participant) + (1|participant: stimulus type) + (1|participant: perspective) + (1|participant: finger) + (1|participant: perspective: hand) + (1|participant: stimulus type: finger) + (1|participant: stimulus type: perspective: hand) + (1|participant: perspective: hand: finger) + (1|participant: stimulus type: perspective: hand: finger).

### Spatial Congruence Analysis

We anticipated that spatial congruence would have important effects on participant performance, and therefore conducted an analysis to specifically examine this effect. Stimuli were grouped according to the spatial congruence between the observed stimulus and required response. Spatial congruence could therefore be either “neutral” (i.e. all middle finger stimuli regardless of perspective, as the middle finger was always in the same central position on screen), “congruent” (i.e. index and ring finger stimuli for the first-person right hand stimuli, and third-person left hand stimuli), or “incongruent” (i.e. index and ring finger stimuli for the first-person left hand stimuli and third-person right hand stimuli). We therefore used a complex random intercept model to see if reaction time was affected by the factors of “Stimulus Type” (2 levels; body or control) and “Spatial Congruence” (3 levels; neutral, congruent, incongruent), using the formula: reaction time ∼ stimulus type * congruence + (1|participant) + (1|participant: stimulus type) + (1|participant: congruence) + (1|participant: stimulus type: congruence).

### Error Rate

The analysis of error rates was carried out in a similar way to the analysis of reaction times, but using binomial generalized linear mixed models to take account of the binary nature of the outcome (0=incorrect, 1=correct). The same general analyses were conducted as for the reaction time. A “Global Analysis” used the formula: error rate ∼ stim_type * perspective * hand * finger + (1 | participant) + (1 | participant: stimulus_type) + (1|participant: perspective: hand)+(1|participant: stimulus type: finger)+(1|participant: stimulus type: perspective: hand: finger), while a “Spatial Congruence Analysis” examined the data using the formula: error rate ∼ stimulus type * congruence + (1|participant) + (1|participant: stimulus type) + (1|participant: congruence) + (1|participant: stimulus type: congruence).

### Response type

We looked at what kind of response participants made when they saw spatially incongruent stimuli. To do this, we first filtered to keep only the spatially incongruent trials. Then the trials were labelled as follows: correct (for trials with a correct response), spatially induced error (for responses given with the index finger when the correct response was the ring finger or vice versa) and spatially neutral error (for incorrect responses given with the middle finger). A generalized linear mixed model was then calculated using the following formula: number of responses ∼ stimulus type * response type + (1|participant) + (1|participant: stimulus type) + (1|participant: response type) + (1|participant: stimulus type: response type).

We were also interested in the reaction time for each type of response. We calculated the following linear mixed model: reaction time ∼ stimulus type * response type + (1|participant) + (1|participant: stimulus type) + (1|participant: response type) + (1|participant: stimulus type: response type).

### Post hoc comparisons

For all models, post hoc comparisons were conducted using z- tests. All p-values obtained were adjusted using Tukey’s method for multiple comparison, and alpha was set to 0.05 for statistical significance.

### Experiment 1 Results

#### Reaction Time

*Global Analysis (fig 2A,B).* Results of the global model are presented in Table 1.

**Figure 2.**
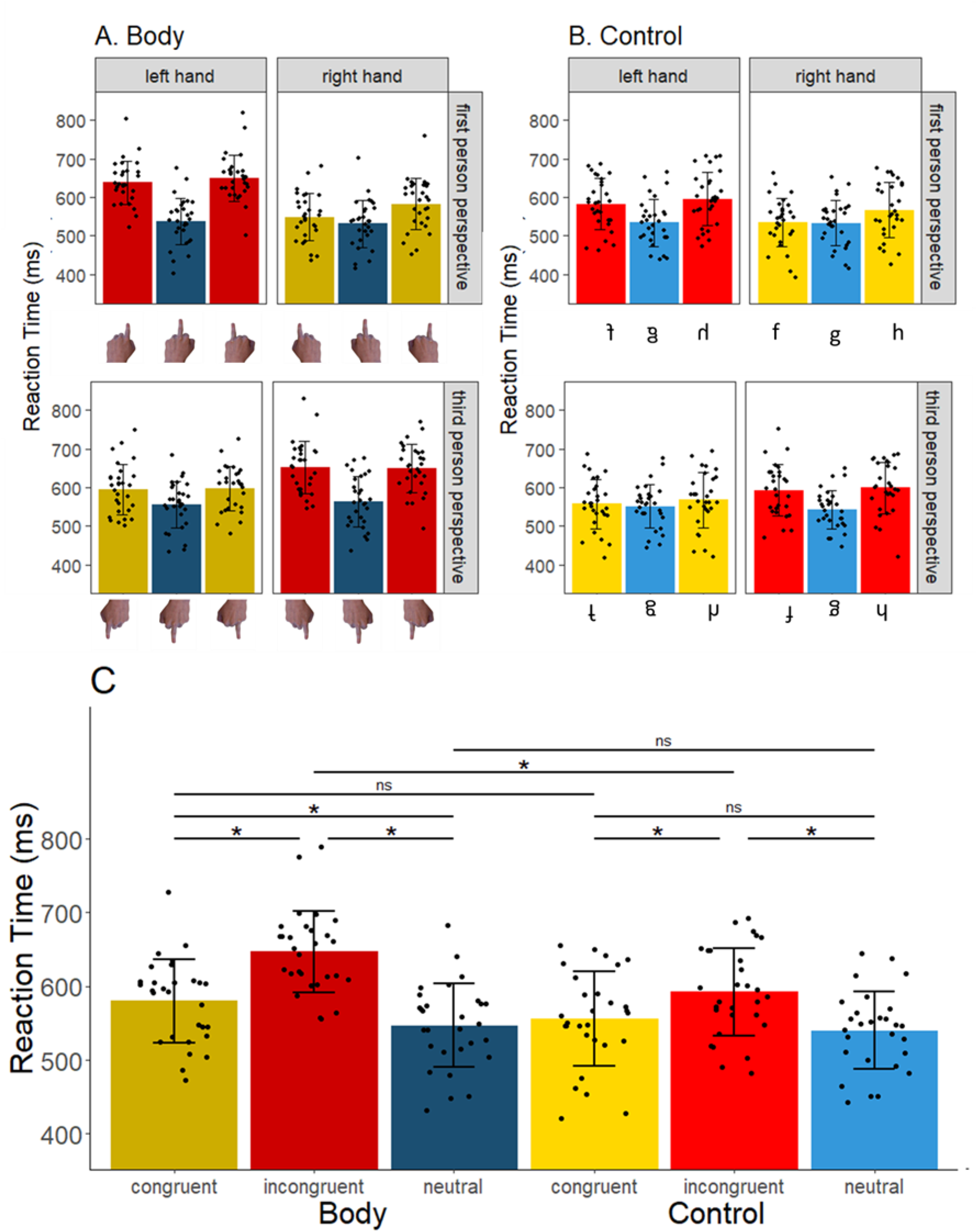
Reaction time analysis. For all plots, yellow bars indicate spatially congruent stimuli, red bars indicate spatially incongruent stimuli, and blue bars indicate spatially neutral stimuli. A darker shade is used for body stimuli and a lighter shade for control stimuli. Error bars represent ±1SD, each point represents a single participant. A) Mean reaction time for all body stimuli. Panels are separated according to hands (columns) and perspective (lines). B) Mean reaction time for all control stimuli. The panels are located in the same relative position as the body stimuli whose characteristics they mimic (see A). C) Mean reaction time for congruent (index or ring finger for the right hand in first-person perspective or the left hand in third-person perspective), incongruent (index or ring finger for the left hand in first-person perspective or the right hand in third-person perspective) and neutral stimuli (all middle finger stimuli).

**Table 1:**
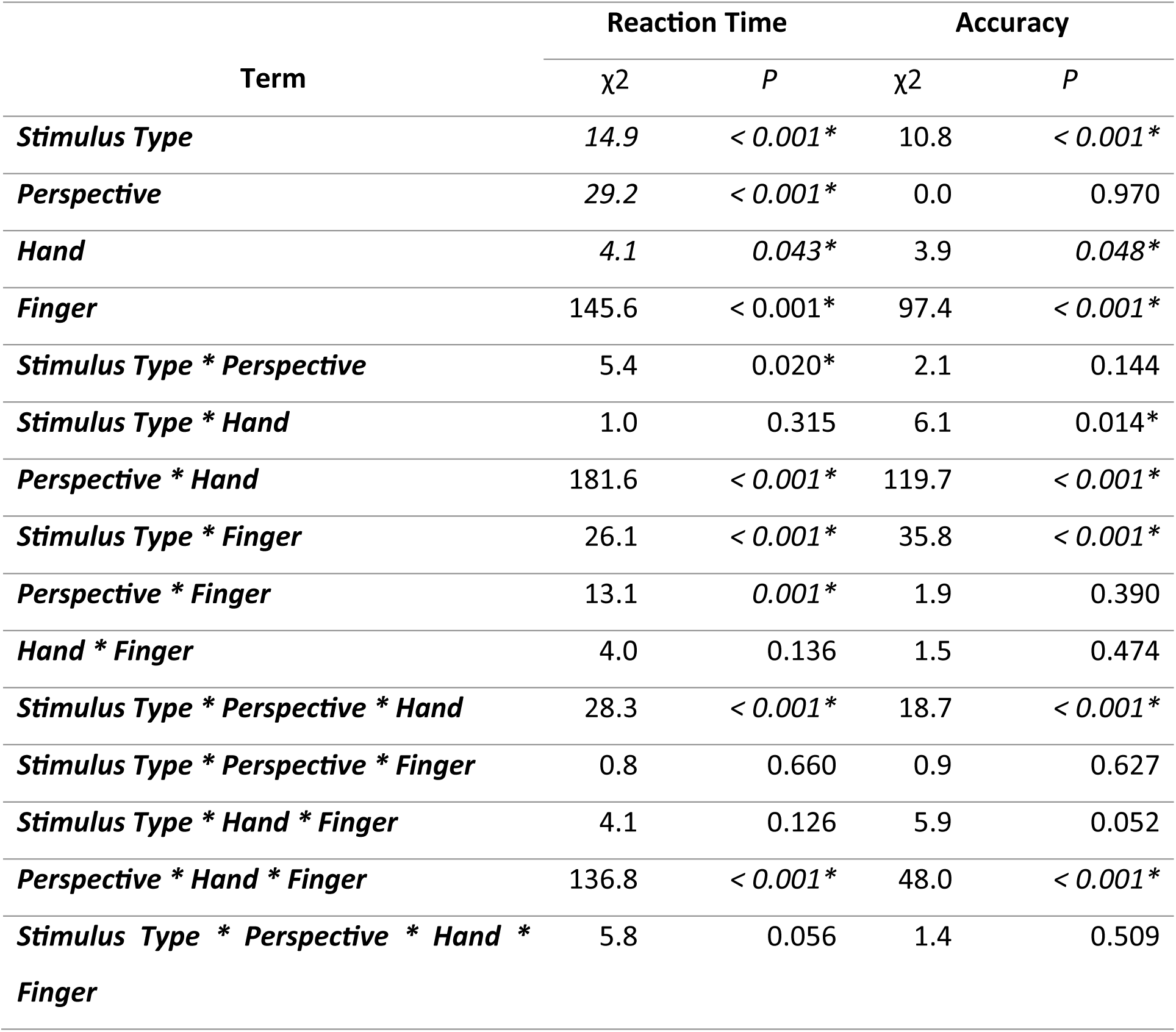
Results from the Global analyses of Reaction Times and Accuracy. * Indicates statistically significant results.

| Term | Reaction Time |  | Accuracy |  |
| --- | --- | --- | --- | --- |
| | $\chi^2$ | *P* | $\chi^2$ | *P* |
| ***Stimulus Type*** | 14.9 | < 0.001* | 10.8 | < 0.001* |
| ***Perspective*** | 29.2 | < 0.001* | 0.0 | 0.970 |
| ***Hand*** | 4.1 | 0.043* | 3.9 | 0.048* |
| ***Finger*** | 145.6 | < 0.001* | 97.4 | < 0.001* |
| ***Stimulus Type \* Perspective*** | 5.4 | 0.020* | 2.1 | 0.144 |
| ***Stimulus Type \* Hand*** | 1.0 | 0.315 | 6.1 | 0.014* |
| ***Perspective \* Hand*** | 181.6 | < 0.001* | 119.7 | < 0.001* |
| ***Stimulus Type \* Finger*** | 26.1 | < 0.001* | 35.8 | < 0.001* |
| ***Perspective \* Finger*** | 13.1 | 0.001* | 1.9 | 0.390 |
| ***Hand \* Finger*** | 4.0 | 0.136 | 1.5 | 0.474 |
| ***Stimulus Type \* Perspective \* Hand*** | 28.3 | < 0.001* | 18.7 | < 0.001* |
| ***Stimulus Type \* Perspective \* Finger*** | 0.8 | 0.660 | 0.9 | 0.627 |
| ***Stimulus Type \* Hand \* Finger*** | 4.1 | 0.126 | 5.9 | 0.052 |
| ***Perspective \* Hand \* Finger*** | 136.8 | < 0.001* | 48.0 | < 0.001* |
| ***Stimulus Type \* Perspective \* Hand \* Finger*** | 5.8 | 0.056 | 1.4 | 0.509 |

The global model identified several notable factors. First, main effects identified that participants were significantly faster when responding to control stimuli (562 ± 57ms) compared to body stimuli (586 ± 53ms, z = 4.0, p< 0.001)(fig S2), when responding to first-person stimuli (566 ± 53ms) compared to third-person stimuli (582 ± 51ms, z = 5.2, p< 0.001)(fig S4), and when responding to right hand (571 ± 51ms) stimuli compared to left hand stimuli (576 ± 51ms, z = 2.1, p = 0.033). This analysis also identified finger-related effects; post-hoc comparisons identified that participants were generally faster when presented with stimuli requiring responses with the middle finger (543 ± 50ms), compared to the index (584 ± 55ms, z = 8.9, p<0.001) or ring fingers (597 ± 54ms, z = 11.6, p<0.001 ), while responses with the index finger were generally faster than those with the ring finger (z = 2.7, p=0.020). Secondly, important interactions between the perspective and observed hand were identified; generally, in the first-person perspective, participants were faster when responding to a right hand (549 ± 55ms) than a left hand (584 ± 52ms) (z = 11.4, p< 0.001), while in the third-person perspective, participants were faster when responding to a left hand (569 ± 51ms) than a right hand (596 ± 51ms) (z = 8.4, p< 0.001). This overall pattern of results was highly consistent with effects that depended on the spatial congruence between the observed stimulus and the required response. As such, further analysis concentrated on examining this effect directly.

### Spatial Congruence Analysis (fig 2C)

The model for spatial congruence showed a significant interaction between stimulus type and spatial congruence (χ^2^ = 39.1, p < 0.001) as well as a main effects of stimulus type (χ^2^ = 15.3, p < 0.001) and spatial congruence (χ^2^ = 266.8, p < 0.001). Post-hoc comparisons focusing on body stimuli showed that reaction time was shortest for neutral (547 ± 57ms) then congruent (580 ± 57ms) then incongruent stimuli (647 ± 55ms) (all z > 5.9, all p < 0.001). Results for the control stimuli showed shorter reaction times for neutral (540 ± 53ms) and congruent stimuli (556 ± 64ms) compared to incongruent stimuli (592 ± 60ms) (both z > 5.9, all p < 0.001). Interestingly, while there was no significant difference in reaction times between neutral body and control stimuli (z= 0.8, p = 0.97) and between congruent body and control participant (z = 2.9, p = 0.05), participants were significantly faster when responding to control stimuli compared to body stimuli spatially incongruent stimuli (z = 6.4, p< 0.001).

### Error rate

#### Global Analysis (fig 3A,B)

Results of the global analysis for error rates is presented in Table 1. Post-hoc comparison showed no significant difference in the error rate between body (10.81 ± 6.11%) and control stimuli (7.30 ± 4.54%)(z = 1.5, p = 0.125). We also identified a finger effect, whereby there were less errors when answering with the middle finger (3.98 ± 3.13%) than with the index (10.06 ± 5.40%, z = 6.1, p< 0.001) and ring fingers (13.16 ± 7.58%, z = 10.4, p< 0.001), whereas the index finger had less errors than the ring finger (z = 4.4, p< 0.001). The analysis also identified several interactions (see Table 1), and the overall pattern of results were again highly consistent with effects depending on spatial congruence. We therefore conducted a specific analysis to examine this effect in greater detail.

**Figure 3.**
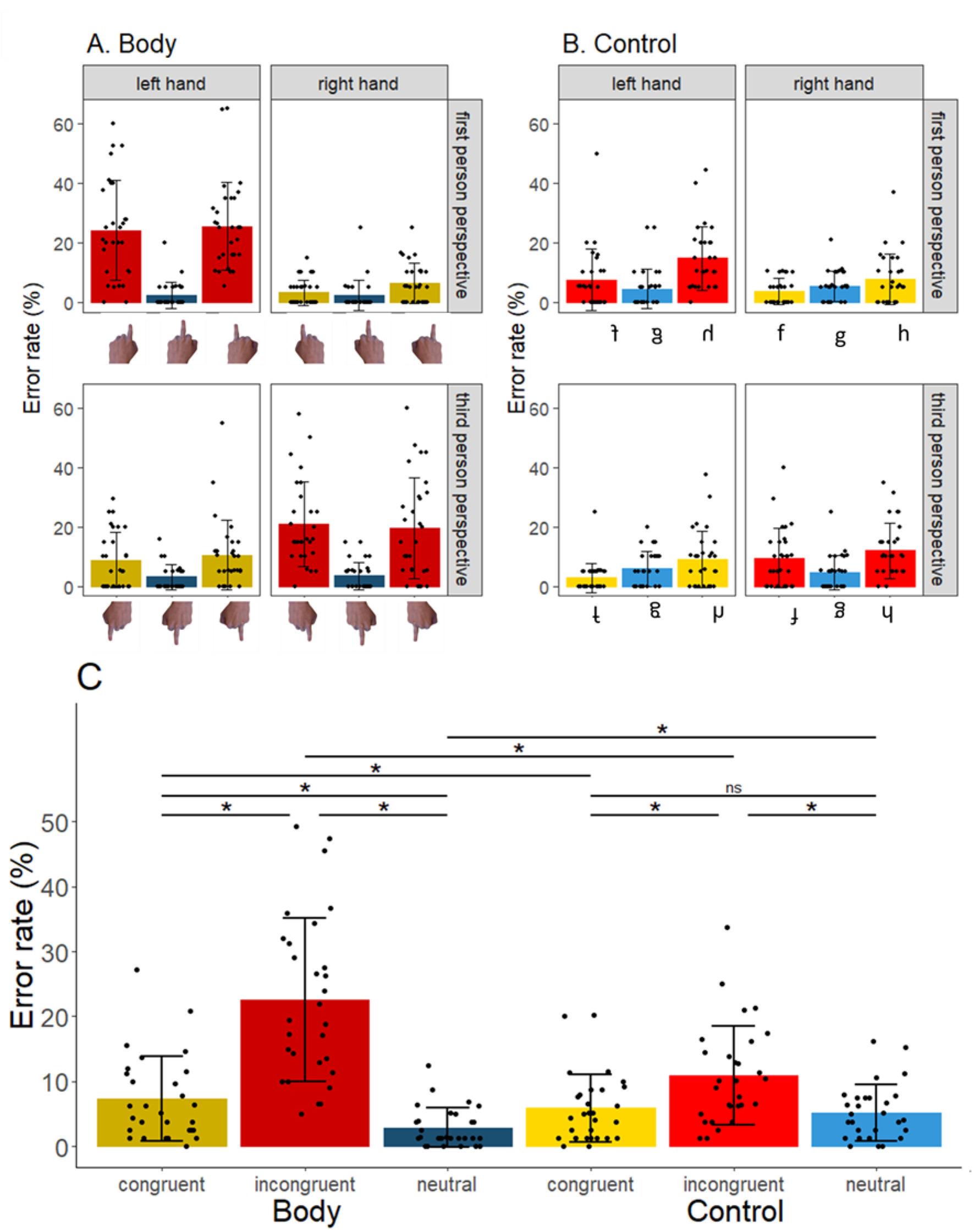
Error rate analysis. Bars are color coded as followed; yellow is spatially congruent stimuli, red is spatially incongruent stimuli, and blue is neutral stimuli. A darker shade is used for body stimuli and a lighter shade for control stimuli. Error bars represent ±1SD, each point represents a single participant. A) Mean error rate for all body stimuli. Panels are separated according to hands (columns) and perspective (lines). B) Mean error for all control stimuli. The panels are located in the same relative position as the body stimuli whose characteristics they mimic (see A). C) Mean error for congruent (index or ring finger for the right hand in first-person perspective or the left hand in third-person perspective), incongruent (index or ring finger for the left hand in first-person perspective or the right hand in third-person perspective) and neutral stimuli (all middle finger stimuli). Body stimuli are on the left and control stimuli on the right.

#### Spatial Congruence Analysis (fig 3C)

The model for spatial congruence showed a significant interaction between stimulus type and spatial congruence (χ^2^ = 40.4, p < 0.001) as well as main effects of stimulus type (χ^2^ = 8.4, p < 0.001) and spatial congruence (χ^2^ = 149.9, p < 0.001). Post-hoc comparisons for the body stimuli showed that the percentage of error was lowest for the neutral (2.85 ± 3.03%), then congruent (7.26 ± 6.53%), then incongruent (22.58 ± 12.63%) stimuli (all z > 4.8, all p < 0.001). Post hoc-analysis for the control stimuli revealed the percentage error did not differ significantly for the neutral (5.13 ± 4.31%) and congruent stimuli (5.84 ± 5.26%) (z = 0.6, p = 0.988), and both were lower than the percentage error for the incongruent stimuli (10.93 ± 7.63%) (both z > 4.3, both p < 0.001).

### Response type Analysis

#### Number of responses (fig 4A)

For this model, the interaction between response type and stimulus type was significant (χ^2^ = 141.7, p < 0.001). The main effect of response type was also significant (χ^2^ = 807.6, p < 0.001) unlike the main effect of stimulus type (χ^2^ = 0.7, p = 0.400). For body stimuli, the number of correct responses (60 ± 10) was higher than the number of spatial errors (15 ± 8, z = 14.1, p<0.001) and the number of ‘spatially neutral’ errors (3 ± 3, z = 21.5, p< 0.001). The number of spatial errors was also higher than the number of ‘spatially neutral’ errors (z = 11.3, p< 0.001). For control stimuli the number of correct responses (70 ± 6) was higher than the number of spatial errors (4 ± 5, z = 22.1, p< 0.001) and the number of ‘spatially neutral’ errors (4 ± 3, z = 22.2, p< 0.001) but there was no difference between the number of spatial and ‘spatially neutral’ errors (z = 0.2, p = 1). All comparisons between corresponding body and control conditions showed less errors for control stimuli (all z > 3.2, all p< 0.018).

**Figure 4.**
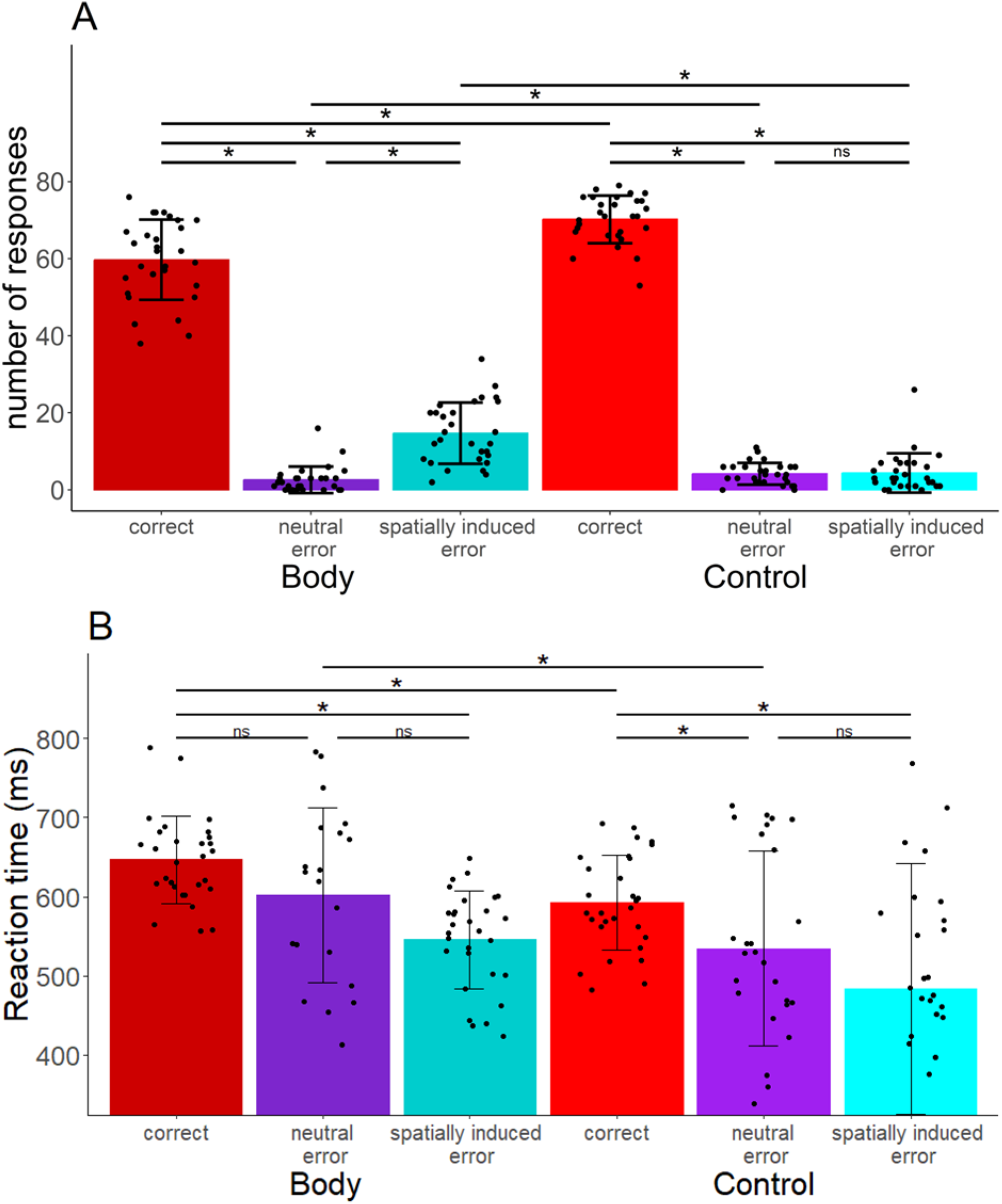
Error type analysis. For all plots, error bars represent SD, each point represents a participant and the color code is as followed; red for correct response, purple for spatially neutral error and cyan for spatially induced error. A darker shade is used for body stimuli and a lighter shade for control stimuli. Error bars represent ±1SD, each point represents a single participant. A) number of response for each type of response for spatially incongruent body and control stimuli. B) Mean reaction time for each type of response for spatially incongruent body and control stimuli.

#### Reaction time (fig 4B)

For the reaction times of these different types of response, the interaction between response type and stimulus type was not significant (χ^2^ = 0.2, p = 0.891). However, the main effect of response type was significant (χ^2^ = 77.4, p < 0.001), as was the main effect of stimulus type (χ^2^ = 34.1, p < 0.001). For body stimuli, the reaction time for correct responses (647 ± 55ms) did not differ to that for ‘spatially neutral’ errors (602 ± 111ms) (z = 2.0, p = 0.337), but was greater than the reaction time for spatial errors (546 ± 62ms) (z = 6.8, p < 0.001). For control stimuli the reaction time for correct responses was the highest (592 ± 60ms) compared to both ‘spatially neutral’ errors (535 ± 122ms) and spatial errors (484 ± 158ms) (both significant z > 3.1, p < 0.022).

### Experiment 1 Discussion

Several noteworthy results emerged from this experiment. Firstly, spatial congruence had a major impact on performance, with longer reaction times and poorer accuracy for spatially incongruent stimuli than for spatially congruent stimuli. Responses to spatially neutral stimuli were fastest with low levels of error, probably because these stimuli were always centred and therefore not subject to spatial congruence effects. Secondly, participants were generally faster and more accurate when responding to control stimuli compared to body stimuli. However, spatial congruence again modulated this effect, as there were no differences in reaction time when participants responded to control and body stimuli presented in a neutral position. Furthermore, our analysis of the errors made indicates that body stimuli were more likely to induce spatial errors, whereas there was no difference between the likelihood of ‘spatial’ and ‘spatially neutral’ errors for control stimuli. These data suggest that the spatial information from body stimuli may be more compelling than for control stimuli. Thirdly, our results indicate that perspective may be less important than would be assumed. Although an effect of perspective was present for reaction time, with faster reaction times for first-person perspective than for third-person perspective, this effect was not present for accuracy. Furthermore, when we break down the results, we see that in first-person perspective the responses are faster and more accurate for stimuli from the (spatially congruent) right hand compared with the left hand (spatially incongruent), whereas for third-person perspective, participants provided faster and more accurate responses to the (spatially congruent) left hand compared to the right hand (spatially incongruent). This again suggests the importance of spatial congruence.

The results of Experiment 1 highlight notable effects of spatial congruence during action observation. They also show a conflict between spatial and anatomical information, suggesting a difference in their processing. This fits well with the hypothesis of two streams of vision: the dorsal stream for processing spatial information and the ventral stream for anatomical recognition (Goodale & Milner, 1992). It should be noted, however, that having two variables of interest (i.e. reaction time and accuracy), can make interpretation of the results complex (e.g the main effect of perspective on reaction time, but not error rates). In addition, the reaction time task gives us information about the ‘end point’ of stimulus processing, but does not give us information about its time-course. Our second experiment addressed these issues using a forced-response paradigm.

## Experiment 2

### Experiment 2 Method

The second experiment was inspired by previous studies that used a ‘forced response’ paradigm to effectively control the time participants have available to prepare a response (Haith et al., 2016; Hardwick et al., 2019; Vleugels et al., 2020) (fig 5A). This allows the measurement of how accuracy changes as a function of the time that participants had to prepare their response. Using this procedure in the present experiment allowed us to assess the amount of time required to process different features of the presented stimuli.

**Figure 5.**
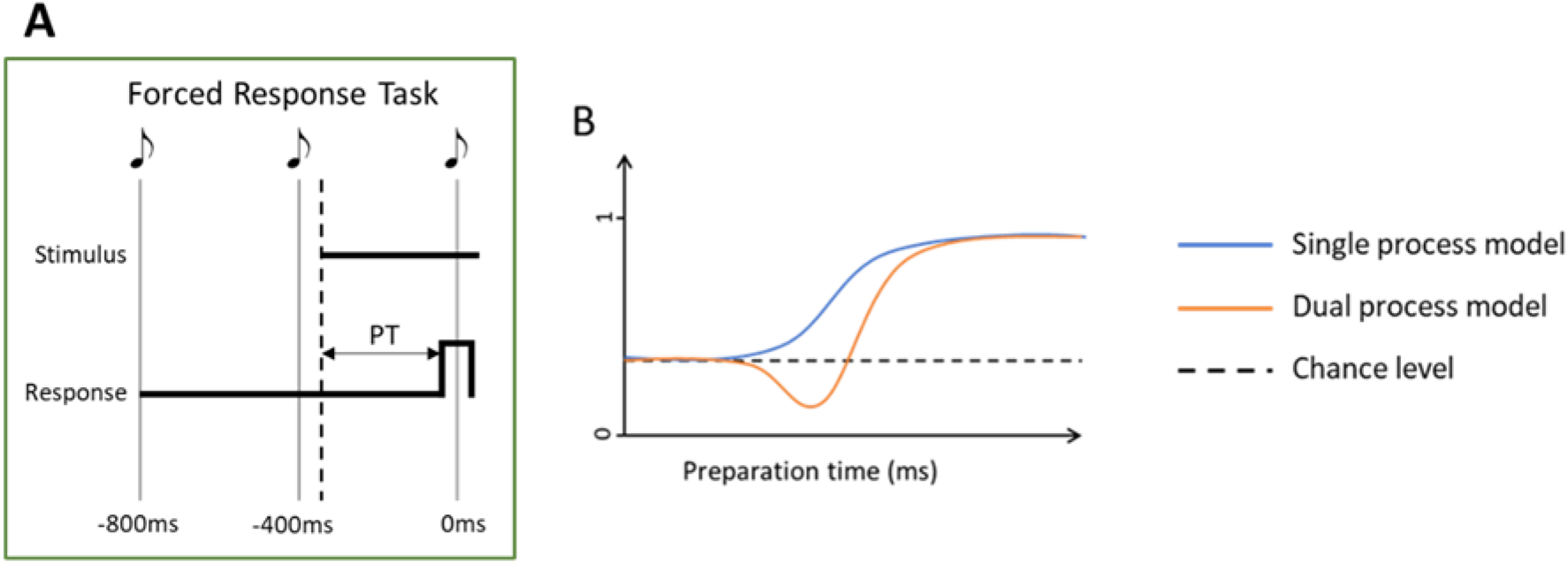
A. Forced Response paradigm. Participants heard a sequence of three equally spaced tones, but here the stimulus appeared at a variable time between the first and third tones. Participants were instructed to respond synchronously with the third tone. This effectively controlled the amount of time that the participant had to process the stimulus and prepare the corresponding response. The preparation time (PT) was defined as the time between the appearance of the stimulus and the participant’s response. B. Models used for analysis of the forced response task. The single process model (blue) follows a sigmoidal response profile, with an initial plateau where the probability of responses begins around chance levels, before increasing to a final plateau where the probability for that response is close to 100%. In orange, the dual process model allows for situations where different responses could be produced as a function of the time available; in the situation shown here an initial decrease below chance level of the probability to produce that type of response before increasing to reach the final plateau.

#### Participants

A group of 57 healthy young participants were recruited via the Prolific online platform (https://www.prolific.com/). Seven participants were excluded from the analyses for failing to follow task instructions (see the exclusion criteria section for more details). This left a total of 50 participants aged between 18 and 35 years (mean = 25.5 years, SD = 4.5 years). All the participants were right handed, 12 were women, 38 were men. Two sample size calculations were performed with Gpower 3.1 based on the results of Hardwick and al., 2019 who used a similar task. The first was based on a period where the two conditions of interest differed by 0.25, based a paired sample, two tailed t-test with alpha = 0.05, power = 0.8 and an effect size (dz) of 2.38. The second followed the same principle but was based on a period with a smaller difference between the two conditions (0.07), with an effect size (dz) of 0.67. The results showed that sample sizes of more than 4 and 20 respectively would be sufficient. To ensure sufficient power in case of additional variability and to ensure that the model fitting would be of high quality, the number of participants was increased to 50. They were all naive about the purpose of the experiment. All participants gave their consent before the start of the experiment and were financially compensated (£5.50). The study was approved by the IPSY ethics committee of UCLouvain.

#### General Task Procedures

The task was created in PsychoPy2 version 2023.2.2 (Peirce et al., 2019) and run online via PsychoPy2’s associated webpage Pavlovia (https://pavlovia.org/) (Bridges et al., 2020).

The general procedure for Experiment 2 was similar to that of Experiment 1. Participants responded to visual stimuli presented in three different conditions; familiarisation, body stimuli, and control stimuli. The stimuli and associated responses were the same as in Experiment 1.

As in Experiment 1, participants heard three short 50ms tones spaced 400ms apart on each trial. Here, however, in each trial a stimulus appeared at a variable time between the first and last tones, and participants were instructed to provide their response synchronously with the third tone (fig 5A). This allowed us to systematically vary the amount of allowed “preparation time” (i.e time between the presentation of the stimulus and the intended time of the response at the third tone; note that all analyses used the ‘actual’ preparation time, i.e. the time between the presentation of the stimulus and the participant’s response) participants had to process the stimulus and prepare a response; presenting the stimulus very early during the sequence of tones allowed a long preparation time, whereas presenting the stimulus very late in the trial provided a very limited preparation time. Note that the timings used included trials in which the stimulus was presented so late that the participant would not realistically have time to process it, forcing them to guess as to which response would be correct. As such, participants were expected to perform at around chance levels when allowed only very limited preparation times, whereas longer preparation times should reflect stimulus processing, allow examination of how response accuracy varied with time. To encourage participants to respond synchronously with the final tone, they were instructed to respond both at the right time and with the correct finger, but that their highest priority was to respond at the right time, even if this meant that they had to guess the correct response. Visual feedback was provided in the same manner as in Experiment 1, and additional feedback was provided to aid participants to respond with good timing. If participants responded 100ms before the third tone, the message "too early" appeared in red on the screen, and if they responded 100ms after the third tone, the message "too late" appeared in red. If they responded between -100ms and +100ms of the third tone, the message "on time" appeared in green. After the third tone there was a period of 800ms during which feedback remained on screen. If the participant provided a response during the trial, there was an interval of 200ms before the next trial. If the participant did not provide a response, the message "no key pressed" appeared on the screen for 2 seconds. As in Experiment 1, between each block, the participants had to rest for at least 10 seconds, during which the instructions were displayed before they could restart. They then had to press the three keys to check that their fingers were correctly in place.

#### Familiarisation Condition

Participants completed 1 block of 144 trials. Each of the three stimuli was presented at the start of the trial (i.e. 0ms, leaving 800ms of preparation time) and then every 17ms thereafter (modelled on the most common 60Hz computer monitor refresh rate) until 799ms (i.e. leaving 1ms of preparation time) for a total of 48 different timings. The order of the trials was pseudo-randomised to ensure that the different timings were distributed throughout the block, and to limit the number of consecutive responses with the same finger to 2. The first two trials were specifically chosen to have longer preparation times to let the participants get used to the task.

#### Body stimuli Condition

Participants completed 4 blocks of 144 trials for a total of 576 trials. The 12 different stimuli each had one trial for each of the 48 timings ranging from 0ms to 799ms before the last tone in steps of 17ms. The trials were distributed over the 4 blocks to ensure an equivalent distribution of each stimulus and each timing. In addition, the order of trials in each block was pseudo-randomised to ensure that the different timings were distributed throughout the block, to limit up to 3 consecutive responses requiring the same button to be pressed, and up to 2 consecutive repeats of the same stimulus.

#### Control stimuli blocks

As with the body stimuli condition, participants completed 4 blocks of 144 trials. The distribution of trials in the different blocks and the pseudo-randomisation were the same as for the body stimuli blocks.

#### Protocol

Participants always began by completing the familiarisation condition, after which condition order was counterbalanced across subjects as in Experiment 1, with half the participants completing the body stimulus condition before the control stimulus condition, and the other half performing the conditions in the opposite order. The median time of completion was 53 minutes, including breaks.

#### Data analysis

Analyses were performed in R version 4.3.2 (R Core Team 2024).

#### Exclusion criteria

In the forced response task, to verify that the participants complied with the instruction to respond at the same time as the third tone, the percentage of responses given within ±100ms of the third tone was calculated for the hand and control blocks. Participants were excluded from our analyses if less than 50% of their responses across the whole experiment were outside the instructed time window. In addition, to ensure that participants were not just concentrating on the timing of their responses without ever trying to answer correctly, participants with a proportion of correct answers of less than chance (33%) throughout the whole experiment, and participants with more than 50% of their answers made using the same finger/key were excluded. As noted above, a total of 7 participants were excluded based on these criteria, leaving a final sample of 50 participants for analysis.

#### Data Visualization

Data were visualized using a sliding window (Haith et al., 2016; Hardwick et al., 2019; Vleugels et al., 2020), considering the proportion of correct/incorrect responses within ±50ms from each time point. We calculated 95% confidence intervals on this data using bootstrapping (1000 iterations) with replacement.

#### Computational Models (fig 5B)

Previous work employing forced response tasks has used computational models to identify different forms of stimulus processing (Haith et al., 2016; Hardwick et al., 2019). The present study compared the ability of two possible models to explain behaviour in the forced response task. These models assume that if participants are forced to respond with very short preparation times, their response will be at random. However, once a participant has sufficient time (t – a random value taken from a gaussian distribution) they will be able to produce a specific response with a high level of accuracy.

The first “single process” model considers only a single process; as such, this model has a sigmoidal form, beginning with responses being given at random, then, as the likelihood that time t has elapsed increases, the proportion of correct responses begins to increase until it reaches a plateau at close to 100% correct responses. This model uses 4 parameters: (1) the initial plateau around responding by chance, (2) the mean and (3) standard deviation of the time t, and (4) the proportion of correct responses at the final plateau. This single process model assumes that there is no conflict between different types of information in the stimulus, allowing the stimulus to be processed as a single global piece of information.

The second “dual process” model assumes that two separate processes take place. This model assumes that after the initial plateau at chance level, an initial, prepotent response becomes available from time t_1_. However, a second response, which requires more processing time to prepare, becomes available at time t_2_, replacing the initially prepared response as the action that will be performed. This would capture phenomenon in which participants would initially prepare a more ‘intuitive’ response (e.g. based on previous experience or salient features of the stimulus), but later replace this with a more reasoned answer. This more complex model includes 7 parameters: (1) the initial plateau around chance levels, (2) the mean and (3) standard deviation of the time t_1_, and (4) the point at which responses based on t1 reach a plateau, (5) the mean and (6) standard deviation of the time t_2_, and (7) a final plateau. This model allows the possibility that there are conflicts between different types of information in the stimulus which are processed in parallel, but require differing amounts of processing time to become available to act upon.

In the present study the data from all the participants was pooled across the relevant condition, and each model was fit to this pooled data. The Akaike information criterion (AIC) was calculated for each model, with comparisons of these scores being used to identify the most favourable model. The code used in the present manuscript was adapted from that used in previous papers using the same model (Haith et al., 2016; Hardwick et al., 2019).

#### SMART method

The SMART (Smoothing Method for Analysis of Response Time-course) method allows investigation of the continuous time-course of the behaviour using cluster-based permutation analysis, providing greater temporal precision when compared to traditional binning methods (for a detailed overview and examples of the code used see (van Leeuwen et al., 2019). First, the time-series of responses was calculated for each participant using a gaussian kernel of width 15ms. The different time-series of the participants were then average weighted across the participants. The 95% confidence intervals were then calculated via the estimated standard error of the weighted mean. Finally, a cluster-based permutation test was performed to determine which cluster was statistically different from a baseline (obtained via the initial plateau parameter of the corresponding model fit described above) or another condition. The code used was that supplied in the original publication describing the procedure (van Leeuwen et al., 2019).

#### Analyses

Results from Experiment 1 indicate that the factor of spatial congruence had important effects on participant performance. Analyses presented in Experiment 2 therefore focus primarily on this factor (though note further analysis of corresponding main effects and interactions are presented in the appendices).

#### Stimulus Type and Spatial Congruence

To examine this interaction, data were categorized into one of 6 conditions based on a 2x3 design according to their stimulus type (i.e. body or control) and spatial congruency (i.e. spatially congruent, incongruent and neutral). Computational models were fit to each condition to establish the complexity of stimulus processing, after which the SMART method was used to identify regions where participant performance differed from their baseline level. Notably, Experiment 1 established that participants had a preference for responding with the middle finger, presumably due to its spatially neutral nature. Therefore, for each condition we used the model fit to establish the baseline likelihood of a given response (i.e. the initial plateau parameter of the fastest process of the model) as the baseline for comparison using the SMART method. Finally, the SMART method was used to compare data for body and control stimuli across corresponding spatial conditions.

#### Response Type Analysis

Following on from the results obtained in the first experiment, we were interested in the type of responses, and more specifically the types of errors, that participants produced for spatially incongruent stimuli. Specifically, if errors were induced by the spatially incongruent nature of the stimuli, we would expect to see more ‘spatially induced’ errors than spatially neutral types of errors. This analysis provided a further in-depth examination of trials in which spatially incongruent stimuli were presented. Again, data were split into 6 conditions based on a 2x3 design based on their stimulus type (body or control) and the response provided (correct response, spatial error, spatially neutral error). Further analysis steps follow the same procedure as that for analysis of stimulus type and spatial congruence above.

#### Perspective analyses

Results from Experiment 1 indicated that effects of visual perspective were secondary to those resulting from spatial congruence. We therefore conducted a series of analyses to assess the presence or absence of effects of perspective without separating the data in relation to spatial congruence. An “interaction” analysis split the data into 4 conditions based on a 2x2 design, considering possible interactions between perspective (first or third-person) and stimulus type (body, control).

All data and code used for the task and data analysis can be found here.

### Experiment 2 Results

#### Stimulus type and spatial congruence

Visual inspection of the results (fig 6A) shows that at very short (less than 200ms) preparation times, participants do not have time to process the stimulus and respond with approximately chance levels of accuracy. However, this initial plateau varies somewhat according to the type of stimulus; spatially neutral stimuli have a slightly higher initial proportion of correct responses. After this initial period as the allowed preparation time increases, there is an increase in the proportion of correct responses up to a plateau close to 100%. There is, however, a notable exception to this pattern of results; for spatially incongruent body stimuli, as the allowed preparation time increases, we initially observe a decrease in the proportion of correct responses, before the increase.

This difference is confirmed by the model fitting results, where the simpler, single process model was able to better explain the data for the majority of conditions (see Table 2); however, data from body, spatially incongruent stimuli were best explained by the dual process model, which could account for a decrease in accuracy below chance levels of performance.

**Table 2:**
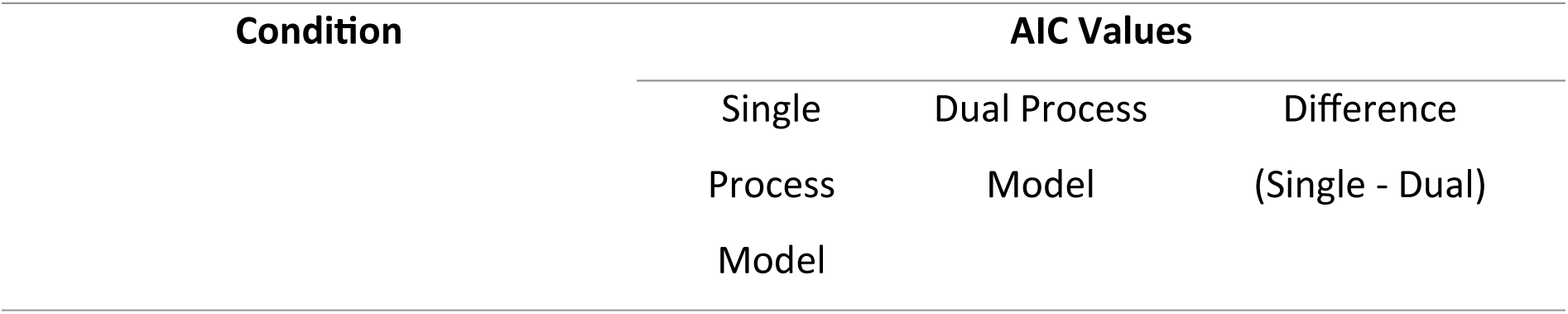
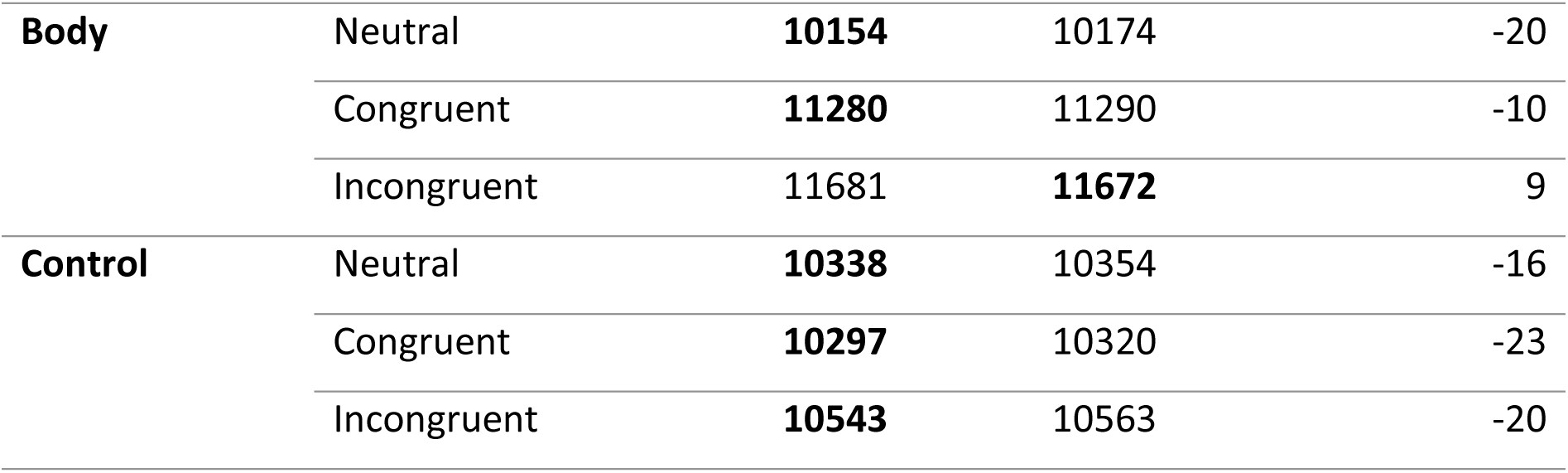
AIC (Akaike Information Criterion) (Akaike, 1974) values for models fit to data from each condition. Bold text indicates the lower AIC score, indicating the better fit model. For difference scores, negative values indicate a better fit for the single process model, and positive values indicate a better fit for the dual process model.

| Condition | AIC Values |  |  |
| --- | --- | --- | --- |
|  | Single<br>Process<br>Model | Dual Process<br>Model | Difference<br>(Single - Dual) |

Table 2: AIC (Akaike Information Criterion) (Akaike, 1974) values for models fit to data from each condition.
|  |  |  |  |  |
| --- | --- | --- | --- | --- |
| **Body** | Neutral | **10154** | 10174 | -20 |
|  | Congruent | **11280** | 11290 | -10 |
|  | Incongruent | 11681 | **11672** | 9 |
| **Control** | Neutral | **10338** | 10354 | -16 |
|  | Congruent | **10297** | 10320 | -23 |
|  | Incongruent | **10543** | 10563 | -20 |

We then used the SMART method to determine detailed comparisons of the time courses for each condition. In the majority of conditions, this analysis demonstrated a relationship whereby accuracy increased with the amount of time participants had to prepare their responses (for full results see Table S2). However, consistent with the computational modelling results, the SMART analysis confirmed that accuracy for body, spatially incongruent data first fell significantly below initial baseline levels from 341-458ms, later rising above the baseline from 500ms onwards.

We then made comparisons between corresponding conditions to identify whether there were any clusters during which the proportion of correct responses was different (fig 6B; for full results see Table S3). For spatially congruent stimuli, a cluster from 394ms indicated a lower final plateau in accuracy for the body stimuli. For spatially incongruent stimuli, a cluster from 334ms indicated that body stimuli showed a lower final plateau and a decrease below the baseline. Finally, we found no significant differences for spatially neutral stimuli presenting body or control images.

#### Response Type Analysis

When visually inspecting the data (Figure 7), we see that the line presenting the likelihood of producing a correct response is the same as the data for the spatially incongruent condition in the “stimulus type and spatial congruence analysis” (i.e. red lines in figures 6 and 7), with a decrease in the proportion of correct responses below chance level only for body stimuli. Visual inspection of the body spatial error shows us that at the same time as the decrease in proportion of correct responses, there is an increase in the proportion of spatial errors (fig 7). Although also present for the control stimuli, this effect is much smaller. The ‘spatially neutral errors’ show the same pattern for the body and control stimuli, with a period at chance level before decreasing to a plateau close to 0%.

**Figure 6.**
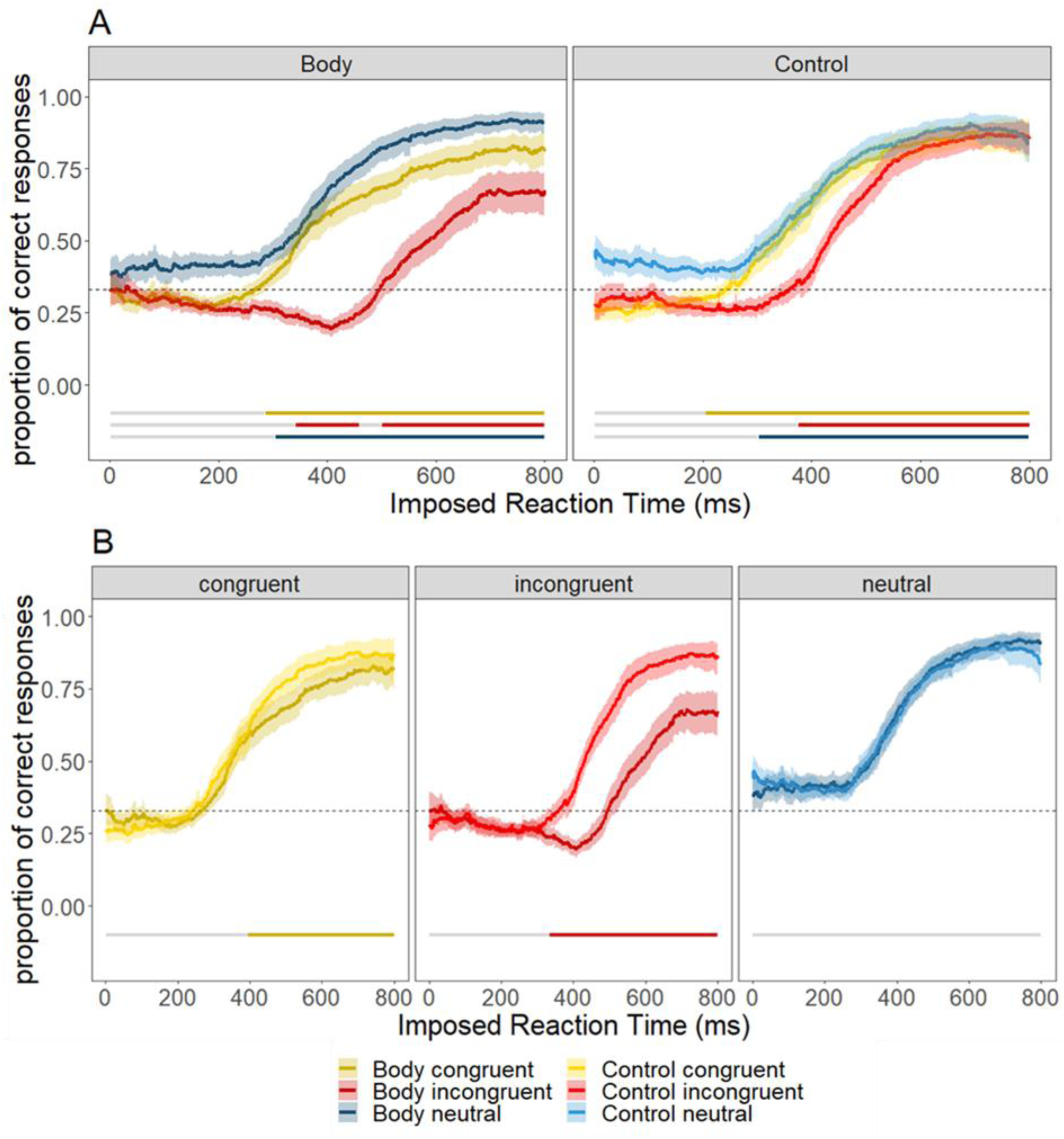
Sliding windows presenting the interaction between stimulus type and spatial congruence. The color code is as followed; yellow is spatially congruent stimuli, red is spatially incongruent stimuli, and blue is neutral stimuli. A darker shade of these colours is used for body stimuli and a lighter shade for control stimuli. Shaded areas present 95% confidence intervals. Lines at the bottoms of the plot indicate significant differences identified using the SMART method, either against baseline (A) or against another condition (B). A. Comparison against baseline. Evolution of the proportion of correct responses as a function of preparation time. The left panel is for body stimuli and the right is for control stimuli. B. Comparison between body and control conditions.

**Figure 7.**
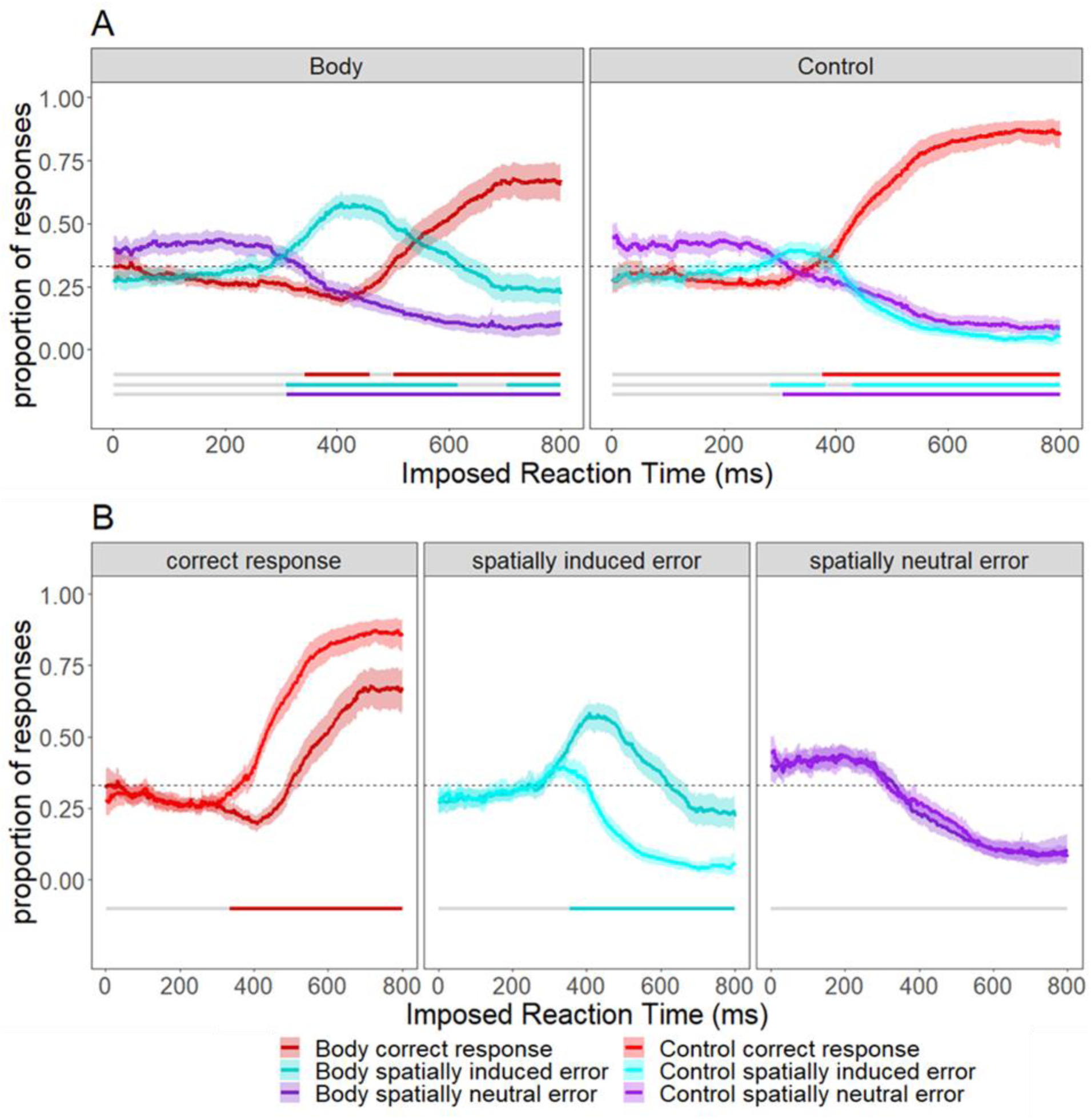
Sliding windows presenting the interaction between stimulus type and type of responses to spatially incongruent stimuli. The color code is as followed; red for correct response, purple for spatially neutral error, and cyan for spatial error. A darker shade of these colours is used for body stimuli and a lighter shade for control stimuli. Shaded areas present 95% confidence intervals. Lines at the bottoms of the plot indicate significant differences identified using the SMART method, either against baseline (A) or against another condition (B). A. Comparison against baseline. Evolution of the probability of a given type of responses as a function of preparation time. B. Comparison between body and control for each type of response.

**Figure 8.**
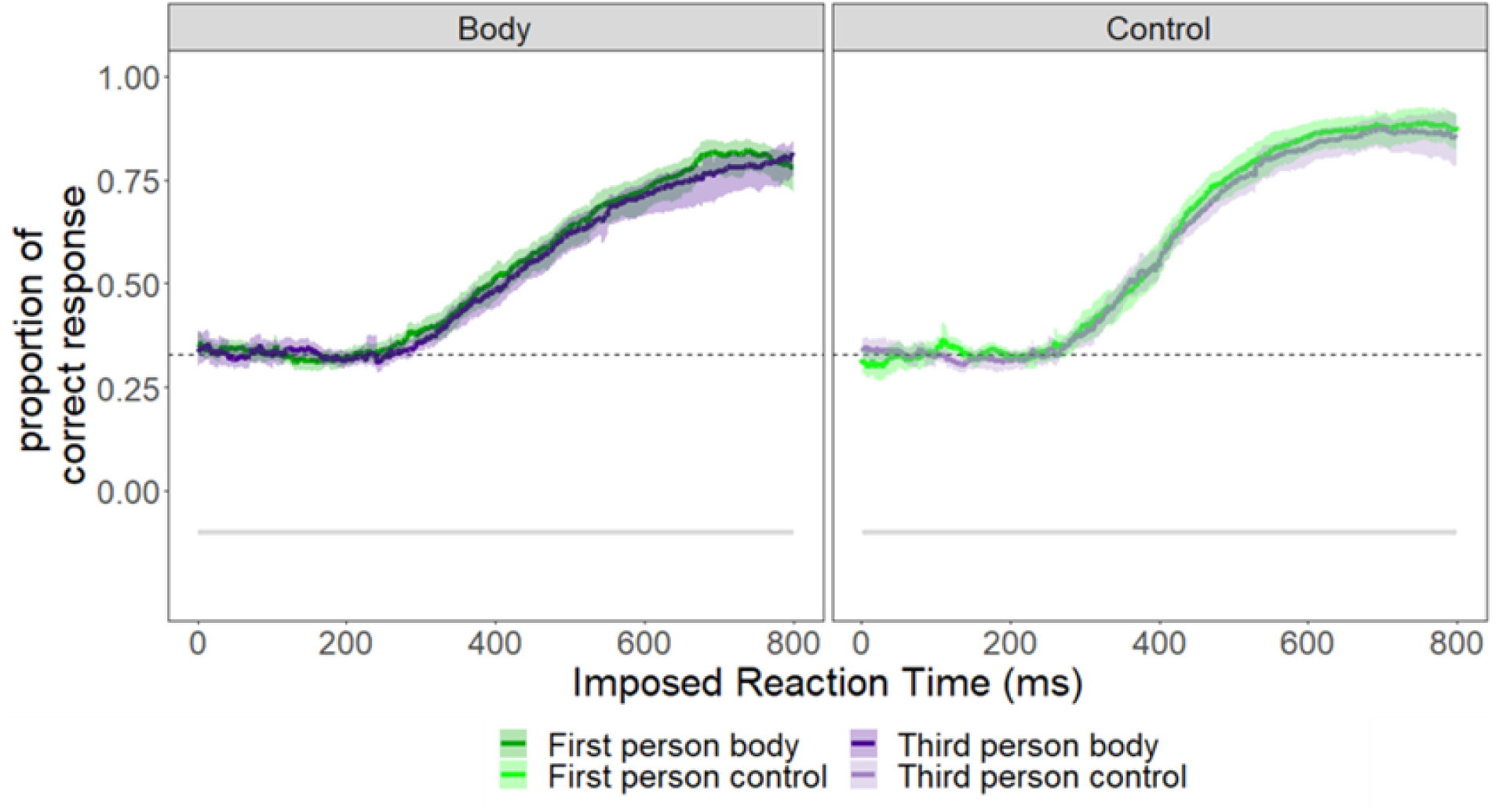
Sliding window visualizations of the data in relation to visual perspective for body and control stimuli. In beige is the evolution for first-person perspective stimuli. In green for third-person perspective stimuli. The lines at the bottom of the plot are coloured when there is a statistical difference between the 2 conditions; note here there were no significant differences across the whole timescale measured for either type of stimuli.

These observations are confirmed by the model fitting results (Table 3). For body stimuli, both correct responses and spatial errors are best explained by the dual process model, whereas spatially neutral responses are best explained by the single process model. For control stimuli, only spatial errors were better explained by the dual process model.

**Table 3:** AIC values for models fit to data from each condition. Bold text indicates the lower AIC score, indicating the better fit model. For difference scores, negative values indicate a better fit for the single process model, and positive values indicate a better fit for the dual process model.

| Response Type |  | AIC Values |  |  |
| --- | --- | --- | --- | --- |
|  |  | Single Process Model | Dual Process Model | Difference (Single - Dual) |
| **Body** | correct | 11681 | **11672** | 9 |
|  | spatial | 12309 | **11991** | 318 |
|  | Spatially neutral | **9585** | 9609 | -24 |
| **Control** | correct | **10543** | 10563 | -20 |
|  | spatial | 8623 | **8609** | 14 |
|  | Spatially neutral | **9864** | 9892 | -28 |

When applying the SMART method (table S2), starting with body stimuli, correct responses first dipped significantly below baseline from 341 to 458ms, before rising significantly above baseline from 500ms onward. By contrast, spatial errors rose significantly above baseline from 307ms to 615ms, before falling significantly below baseline from 730ms onwards. In comparison, spatially neutral errors followed a simpler trajectory, dropping significantly below baseline from 310ms onwards, and remaining low thereafter. For the control stimuli, in contrast to the body stimuli, correct responses never fell below baseline levels, and instead increasingly only significantly above baseline from 374ms. By contrast, spatial errors rose significantly above baseline from 282ms to 383ms, before falling significantly below baseline levels from 429ms. Finally, spatially neutral errors fell significantly below baseline from 303ms.

We then made comparisons between corresponding conditions (fig 7B)(table S3). For correct responses, while the curves were similar, body responses fell significantly below control stimuli from 334ms onwards, with a slower inflection and a lower final plateau. We found a corresponding effect on spatial errors; again, the curves had similar shapes, but differed from 355ms onwards, with a greater likelihood of errors for body compared to control stimuli. Finally, comparison of the curves for spatially neutral errors found no differences between the body and control conditions.

#### Perspective analysis

Visual inspection of the data indicated that all data followed a generally sigmoidal shape, with accuracy generally rising more slowly for body than control stimuli. Model fitting identified that for all conditions, the single process model was the most favourable (all AIC diff < - 17)(table S1-2). Using the SMART method, we found that the proportion of correct responses increased significantly above baseline for all conditions (see Table S3), though the rise occurred more slowly for body compared to control stimuli. Notably, when using the SMART method to directly examine possible effects of perspective, we found no significant differences between first- and third-person stimuli for the body condition, nor were there any differences when comparing the corresponding stimuli in the control condition.

### Experiment 2 Discussion

Several noteworthy results can be highlighted in this experiment. Firstly, as in Experiment 1, we can see a major effect of spatial congruence with, in particular, a difference in behaviour for spatially incongruent body stimuli. Indeed, these stimuli are better explained by a dual-process model, which can account for an initial decrease in the proportion of correct responses before an increase. These results are consistent with participants making incorrect responses based on *where* the stimulus is presented, rather than *what* it presents. This is supported by the type of response participants made to spatially incongruent stimuli; as the proportion of correct responses decreases, there is a corresponding increase in spatial errors. Interestingly, this pattern of results was markedly stronger for body stimuli. Indeed, for control stimuli, there was no decrease in the proportion of correct responses, and the increase in spatially induced responses, although significant, was much smaller than for the body stimuli. These results, which are in line with those of Experiment 1, therefore show a difference between the processing of body and control stimuli, and more particularly of spatial information. Finally, while Experiment 1 identified a weak effect of perspective, in Experiment 2 there were no significant differences between first- and third-person perspective for either body or control stimuli. This result again emphasizes that the importance of perspective during action observation may be overestimated when compared to spatial congruence.

## General Discussion

The present study characterized the role of the factors of stimulus type (body or control), spatial congruence, and perspective (first- and third-person) during action observation. Experiment 1 used a classical reaction time paradigm to examine how participants responded to these factors. Experiment 2 used a forced response paradigm to examine the time-course of stimulus processing, and how the effects of these factors varied depending on the allowed processing time.

Results across the two experiments indicate that spatial congruence has a major impact on the observation of actions, as introducing conflicts between the required response and the spatial location of the stimulus led to a significant increase in incorrect responses. Interestingly, this effect was stronger in body compared to control images, consistent with a difference in the processing of these two types of stimuli. Furthermore, in contrast with previous reports, we found little difference for first-person perspective over third-person perspective, and our results suggest such effects could be mostly driven by spatial congruence.

### Stronger spatial congruence effects for body stimuli

The present experiments indicate that the spatial congruence of the stimulus has a major impact on participants’ performance. Interestingly, this effect was more pronounced for body- related stimuli than for control stimuli. In particular, results of Experiment 2 indicate that participants initially respond on the basis of the stimulus’ spatial information, even if this is irrelevant for this task, before this initial automatic response is replaced by a response based on anatomical recognition. It therefore seems that there are different timescales of information processing for spatial and anatomical information. Interestingly, these effects were smaller or absent for control stimuli, consistent with previous work proposing differences in their processing (for review see (Gowen & Poliakoff, 2012)).

These results are consistent with the ‘dual visual streams hypothesis’ (Goodale & Milner, 1992), which proposes that spatial processing (’where’) and object recognition (‘what’) are respectively processed by separate dorsal and ventral visual steams. In particular, the dorsal stream is known for its role in visually guided actions, which require rapid visual processing to allow us to quickly adapt movements according to visual feedback (Króliczak et al., 2008). By contrast, the ventral stream processes information related to visual form, and does not have the same level of immediacy. Thus, in the context of Experiment 2, which examined the time- course of information processing, spatial information processed via the dorsal stream would be available to influence participant responses earlier than anatomical information processed via the ventral stream. The differences observed between the body and control stimuli could be explained by three non-exclusive proposals. The first is that the dorsal stream is particularly sensitive to body-related stimuli. Motor simulation theory proposes that action observation activates the same neural circuitry as action execution (Jeannerod, 2001), and since the dorsal stream is known for its role in visuomotor control (Goodale & Milner, 1992), it is possible that it is more strongly attuned to body stimuli. In addition, the dorsal stream contains areas such as the inferior parietal lobule (Freud et al., 2016), known to contain mirror neurons that are activated during observation of action, and thus in the present experiment would be recruited only by our body stimuli. Increased activation of these regions by observation of body stimuli could in turn make the information processed by the dorsal stream more compelling. The second is that the dorsal stream passes information to premotor areas that are activated during action observation (Goodale & Milner, 1992), and this activation could be necessary to induce this response based on spatial information. The third is that the differences could be explained by the choice of control stimuli (see strengths and limitations section for more details on the choice of stimuli). In fact, for control stimuli, the letter is represented on its own without being linked to any other reference element, which is not the case for body stimuli where other information is present (i.e. the hand). These differences between the stimuli could also partly explain the results obtained.

### Effect of perspective

The results suggest that perspective has a relatively minor effect on action priming, and that effects of perspective may in fact mostly be driven by spatial congruence. Effects of perspective in Experiment 1 were limited only to reaction times, which were faster for first- than third-person perspective, with differences that were smaller in magnitude than those identified for spatial congruence. Similarly, no effects of perspective were identified in Experiment 2. These results suggest the effects of perspective described in the literature so far could be partially or even mostly driven by spatial congruence. Indeed, several studies on perspective have compared right hands in first-person perspective with right hands in third-person perspective (Angelini et al., 2021; Ge et al., 2018; Oosterhof et al., 2012);our present results suggest that the differences observed in such experiments may be driven more by a change in spatial congruence than by a change in perspective. This interpretation is in agreement with other results (Avikainen et al., 2003; Bekkering et al., 2000; Chiavarino et al., 2007; Shmuelof & Zohary, 2008; Wohlschläger & Bekkering, 2002) that show an interaction between perspective and the identity of the hand observed. It therefore seems important that future studies on the effects of perspective during action observation choose their stimuli carefully. An optimal solution would be to compare both right and left hands in first- and third-person perspective, but in cases where the overall number of trials needs to be limited, comparing a right hand in a first-person perspective with a left hand in a third-person perspective would allow ‘true’ comparisons between perspectives that are not potentially confounded by differences in spatial congruence.

### Effect of stimulus type

Results of the present study indicate that participants were generally faster and more accurate when responding to control stimuli compared to body stimuli. We note, however, that the main effect of stimulus type was in fact likely driven by the interaction between stimulus type and spatial congruence, as we identified minimal differences in responses to spatially neutral body and control stimuli in Experiment 1, and no differences in Experiment 2. These results go against the current literature, which shows a tendency towards a preference for body stimuli, or a status quo between the two (Gowen & Poliakoff, 2012). One possible explanation is that, as proposed by Gowen and Poliakoff, the priming effect induced by different types of stimuli depends mainly on their characteristics (e.g. saliency, size) and not on whether they are “body-related” in nature. It is therefore possible that the low-level properties of the stimuli were one of the causes of the preference for control stimuli. Specifically, even though the position of the relevant part of the stimulus to respond to (i.e. the letter vs the finger) was presented in a consistent manner across our body and control stimuli, the body stimuli presented more overall information (i.e. not only the relevant finger, but also the rest of the hand) than the control stimuli (which showed only the relevant letter on an otherwise blank screen). We note that the selection of control stimuli for action observation studies is a complex issue, (see for example (Loporto et al., 2011)) and provide further discussion of this point in the ‘strengths and limitations’ section below. (Holahan et al., 1978)

### Strengths and limitations

One of the main strengths of this paper is the information provided by the forced response task. Without it, we would not have been able to observe the differences in information processing, particularly between spatially incongruent body and control stimuli. The use of this paradigm therefore enabled us to obtain important new temporal information. In addition, analysis using the SMART method meant that we did not lose the precision of the temporal information that would have been the case with binning analysis (van Leeuwen et al., 2019).

The choice of suitable control stimuli is a difficult one as there are an infinite number of possibilities. The approach used in the present study allowed us to maintain the position of the most relevant information of interest across equivalent stimuli. This approach appears to have been somewhat successful, as responses to spatially neutral stimuli did not differ for any of our comparisons between body-related and control stimuli. However, it is not possible to generalise our results to all possible control stimuli. Certain parameters, such as the salience of the stimuli, may not have been perfectly matched up with their body-related counterparts. A limitation of the stimuli we have chosen is the fact that for body stimuli, the fingers are represented in relation to the hand, which is not the case with the control stimuli, where the letters stand alone. It is possible that the difference in the effect of spatial congruence between body and control stimuli is partly influenced by this. Similarly, low-level perceptual properties of the stimuli (e.g. the number of white/coloured pixels) differed between our stimuli. However, we again note that there were no significant differences between spatially neutral stimuli in the body-related and control conditions, suggesting that reaction times were mainly influenced by other factors. In addition, the body stimuli represented raised fingers, while the participants had to lower theirs to respond. This could have the effect of globally increasing the reaction time for the body stimuli, as the stimuli are imitatively incompatible with the movement the participants have to perform. Critically, the limitations outlined above do not undermine our main findings, which highlight an effect of spatial congruence on body-related stimuli, which will still be correct even if the control results are not generalizable. Nonetheless, further studies with different types of control stimuli are needed to test this generalisation.

For the body stimuli, we used static stimuli. There were two main reasons for this choice. Firstly, the aim of the forced response task was to have precise control over the time participants had to prepare a response and process the stimuli. The use of static stimuli allowed more precise timing control than the use of video. Secondly, previous studies have already used static stimuli in the context of action observation. For these reasons, static stimuli seemed the most appropriate. However, future studies are needed to confirm that these results can be generalized to video viewing.

Finally, the results of Experiment 2 suggest that a ‘finger effect’ was present in the forced response task. At very short reaction times (approximately <200ms), if responses were given truly randomly, we would expect the proportion of responses with the index, middle and ring fingers should be the same (i.e. 33%). However, we found that around 40% of responses were with the middle finger and 30% with the index and ring fingers. This preference could be explained by the fact that these stimuli never presented any conflict between spatial and anatomical information for responses with the middle finger. Future studies using similar methodology could ask whether this represents a conscious strategy by interviewing participants during debriefing. This preference has the effect of modifying the baseline when the stimuli requiring a response with the middle finger are separated from the others (thus in the analysis of spatial congruence) with a higher baseline for neutral stimuli than for congruent or incongruent stimuli. The possible impact that this effect could have is minimised (or even eliminated) thanks to our analysis method, which allows us to calculate the baseline specific to each condition, and then compare how the proportion corresponding responses varies in relation to this baseline (note that with SMART method, we only made pairwise comparisons between conditions with similar baselines).

It is important to note that the results of the present study are exclusively at the behavioural level; proposals of the neural mechanisms behind the effects on reaction time and accuracy, while based on results from the previous literature, are therefore speculative at this stage. Future experiments could explore the neural correlates of these effects in further detail, for example, by using repetitive Transcranial Magnetic Stimulation (TMS) to disrupt areas of the dorsal and ventral visual streams, and see how this affects performance in the task. In addition, given the time-dependent nature of the present results, using techniques such as TMS to test how corticospinal excitability is modulated by different stimuli at different timings, or electroencephalography to examine changes in evoked potentials at different stages of stimulus processing, present interesting avenues for future research

## Conclusion

These experiments show that spatial congruence has a major impact on action observation, with longer reaction times and more errors when there is a conflict between the response requested and the spatial position of the stimulus. This phenomenon can be explained by the fact that spatial information is processed more quickly than anatomical information, thus inducing a first automatic response based on spatial information that may be incorrect. Interestingly, this phenomenon is more pronounced for body-related than control stimuli, which is consistent with a difference in processing between these two types of stimuli. Furthermore, we found little difference between first- and third-person perspective. Our results also suggest that the perspective effect previously identified in the literature may be driven mainly by spatial congruence. We believe that a better understanding of how stimuli are processed during action observation is important for future applications. For example, future research into the use of action observation for rehabilitation after stroke may consider optimizing the spatial compatibility of the stimulus and the observer, as this seems to be optimal for participant performance.

## Data availability statement

All data and code used for the task and data analysis can be found here: https://osf.io/q2rd8/?view_only=e492221e800a4d78b44040910e10985b

## Author Contributions (CRediT)

| Author | Conceptualization | Methodology | Software | Validation | Formal Analysis | Investigation | Resources | Data Curation | Writing – original draft | Writing – reviewing & editing | Visualization | Supervision | Project Administration | Funding Acquisition |
| --- | --- | --- | --- | --- | --- | --- | --- | --- | --- | --- | --- | --- | --- | --- |
| Baptiste M. Waltzing | x | x | x | x | x | x | x | x | x | x | x |  |  |  |
| Siobhan McAteer |  |  | x |  | x |  |  |  |  | x |  |  |  |  |
| Marcos Moreno-Verdú |  |  | x |  | x |  |  |  |  | x |  |  |  |  |
| Elise E. Van Caenegem |  |  |  |  |  |  |  |  |  | x |  |  |  |  |
| Yue Du |  |  | x |  |  |  |  |  |  |  |  |  |  |  |
| Robert M. Hardwick | x | x | x | x | x |  | x |  | x | x |  | x | x | x |

## Diversity in Citation Practices

Retrospective analysis of the citations in every article published in this journal from 2010 to 2021 reveals a persistent pattern of gender imbalance: Although the proportions of authorship teams (categorized by estimated gender identification of first author/last author) publishing in the *Journal of Cognitive Neuroscience* (*JoCN*) during this period were M(an)/M = .407, W(oman)/M = .32, M/W = .115, and W/W = .159, the comparable proportions for the articles that these authorship teams cited were M/M = .549, W/M = .257, M/W = .109, and W/W = .085 (Postle and Fulvio, *JoCN*, 34:1, pp. 1–3). Consequently, *JoCN* encourages all authors to consider gender balance explicitly when selecting which articles to cite and gives them the opportunity to report their article’s gender citation balance.

## Acknowledgments

None

## Appendices

### 1. Results Experiment 1

#### 1.1 Speed -accuracy trade-off

A speed-accuracy trade-off was plotted primarily to spot a potential outlier. After visual inspection, one participant was identified as an outlier due to a very high percentage of error (close to 50%) and very short reaction time. This participant was excluded for all analysis. A Pearson correlation coefficient was calculated. It was significant (r(28) = -0.42, p = 0.02) before the exclusion of the participant but didn’t hold after the exclusion (r(28) = -0.17, p = 0.37).

**Figure S1.**
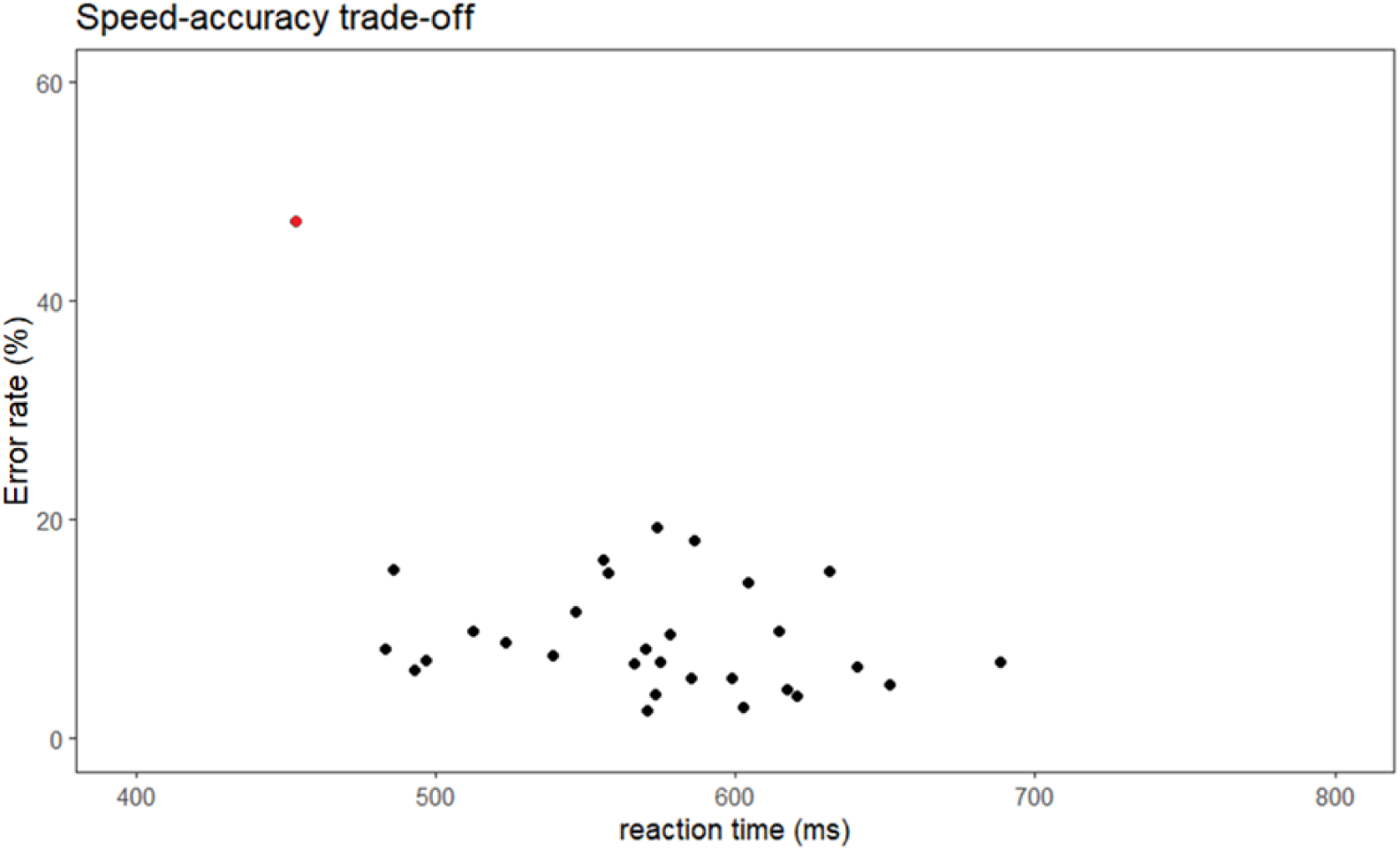
Speed-accuracy trade off. Each point represents a participant. The red point is the excluded participant.

#### 1.2 Stimulus type

**Figure S2.**
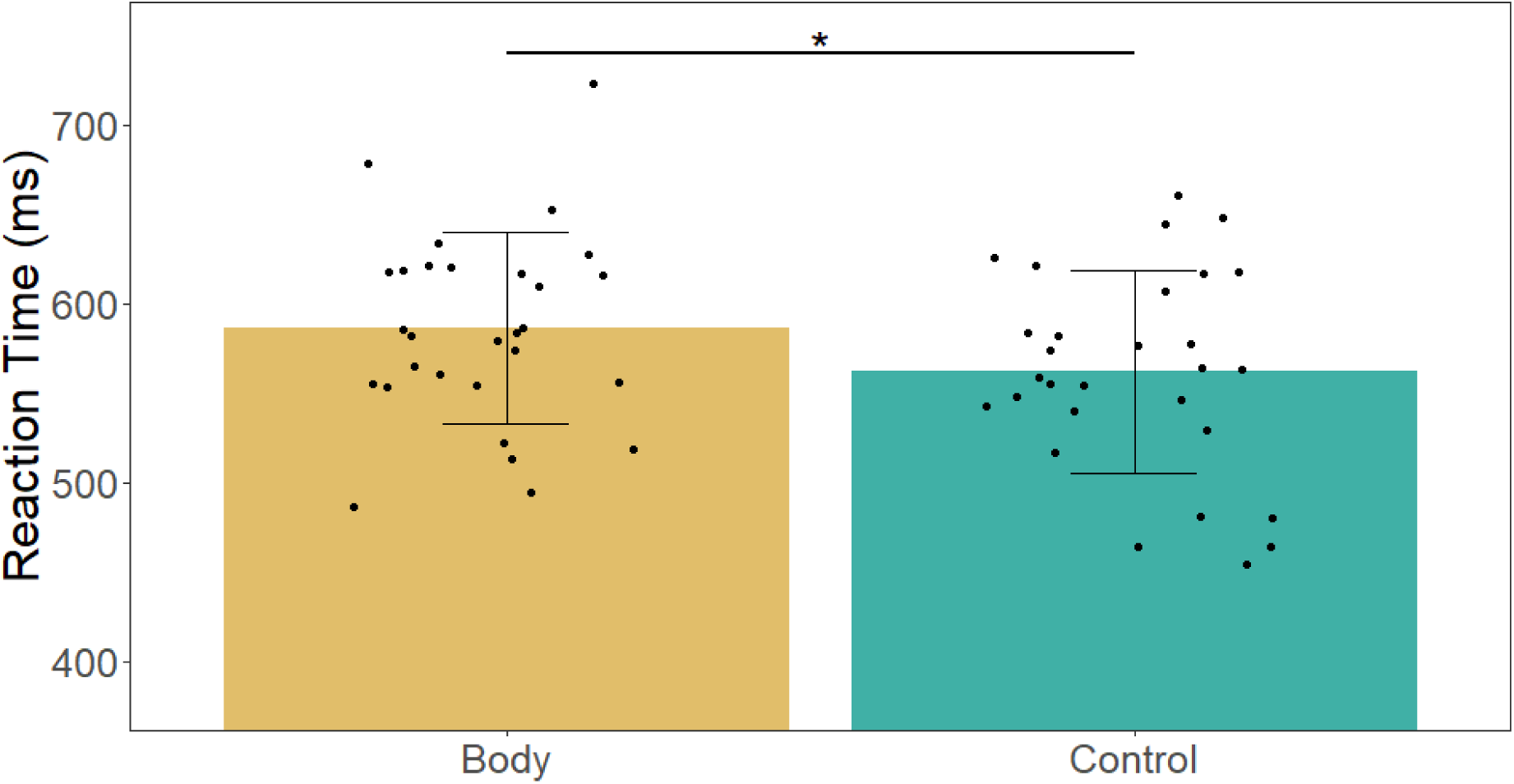
Reaction time for body and control stimuli. Error bars represent SD, each point represents a participant.

**Figure S3.**
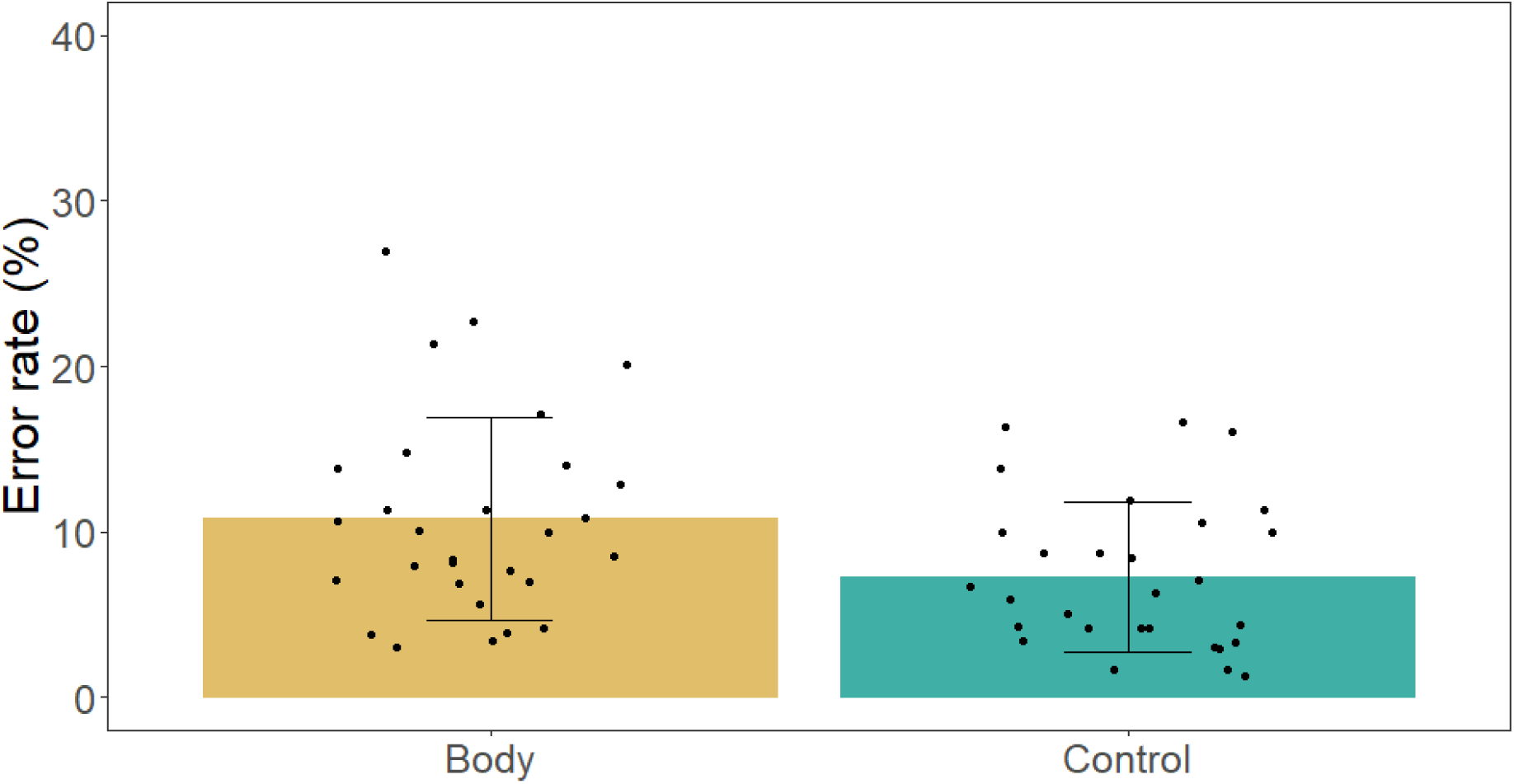
Error rate for body and control stimuli. Error bars represent SD, each point represents a participant.

#### 1.3 Perspective

**Figure S4.**
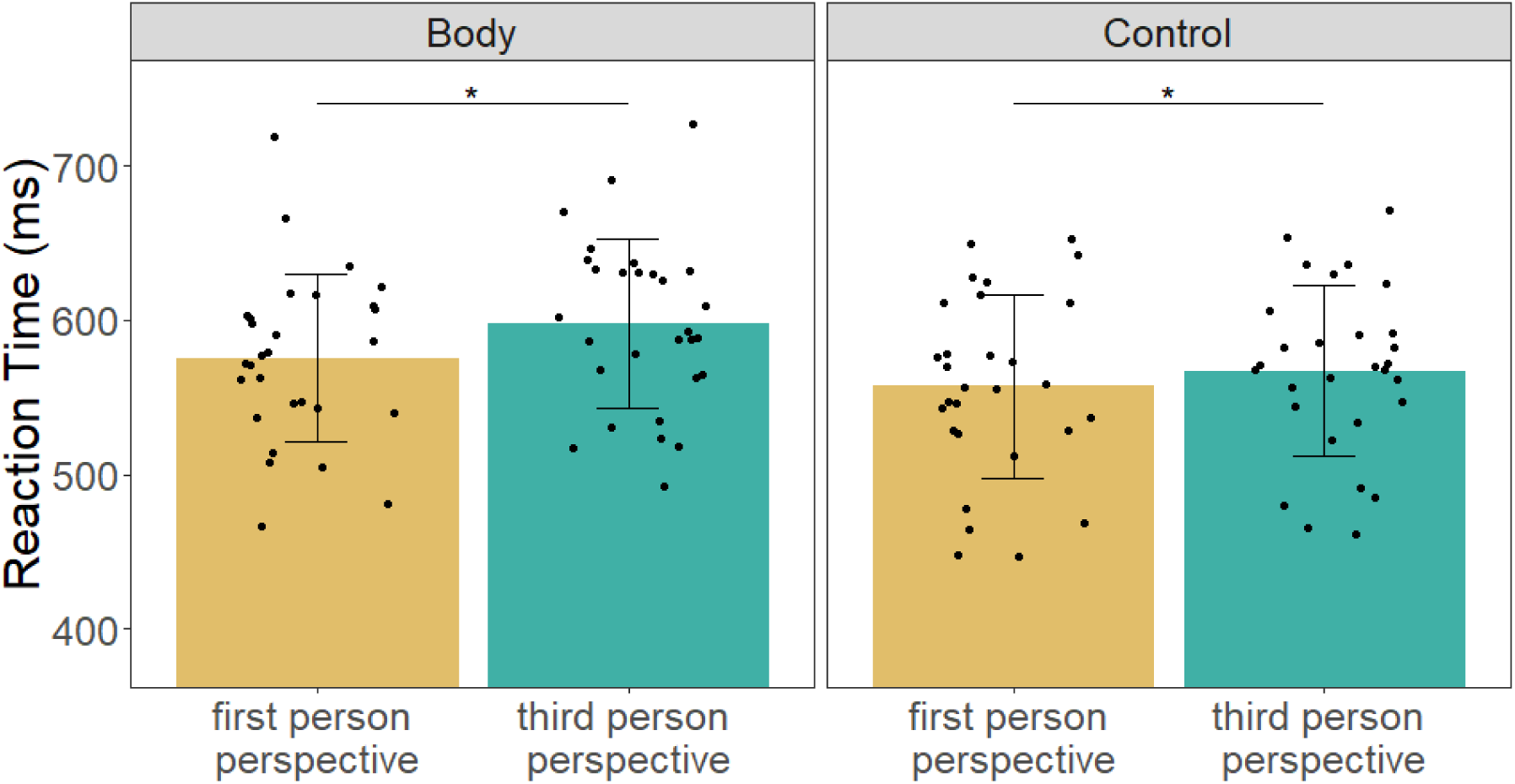
Mean reaction time for stimuli in first-person perspective and third-person perspective. The left panel is for body stimuli and the right is for control stimuli. Error bars represent SD, each point represents a participant.

**Figure S5.**
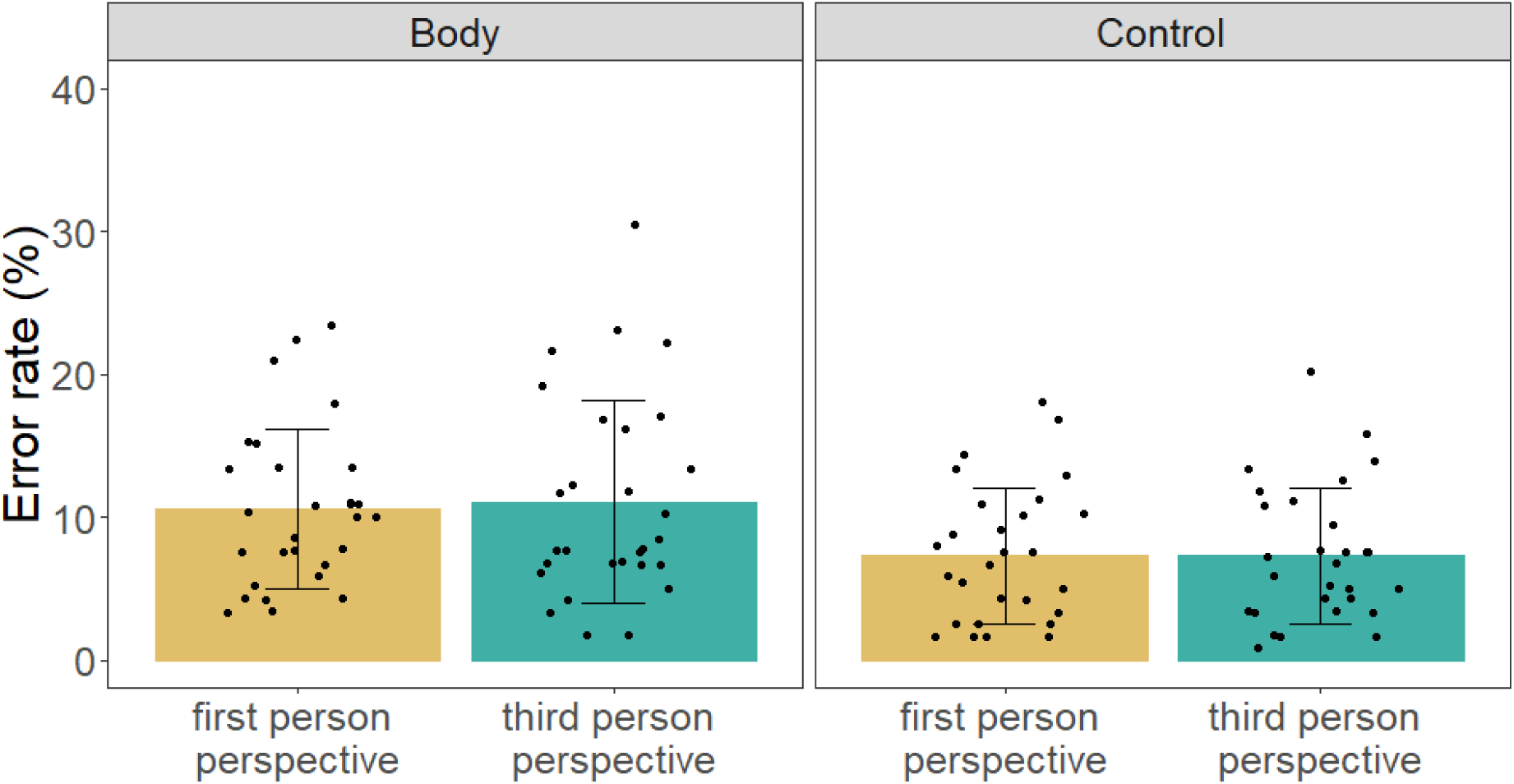
Mean reaction time for stimuli in first-person perspective and third-person perspective. The left panel is for body stimuli and the right is for control stimuli. Error bars represent SD, each point represents a participant.

### 2. Results Experiment 2

**Table S1.**
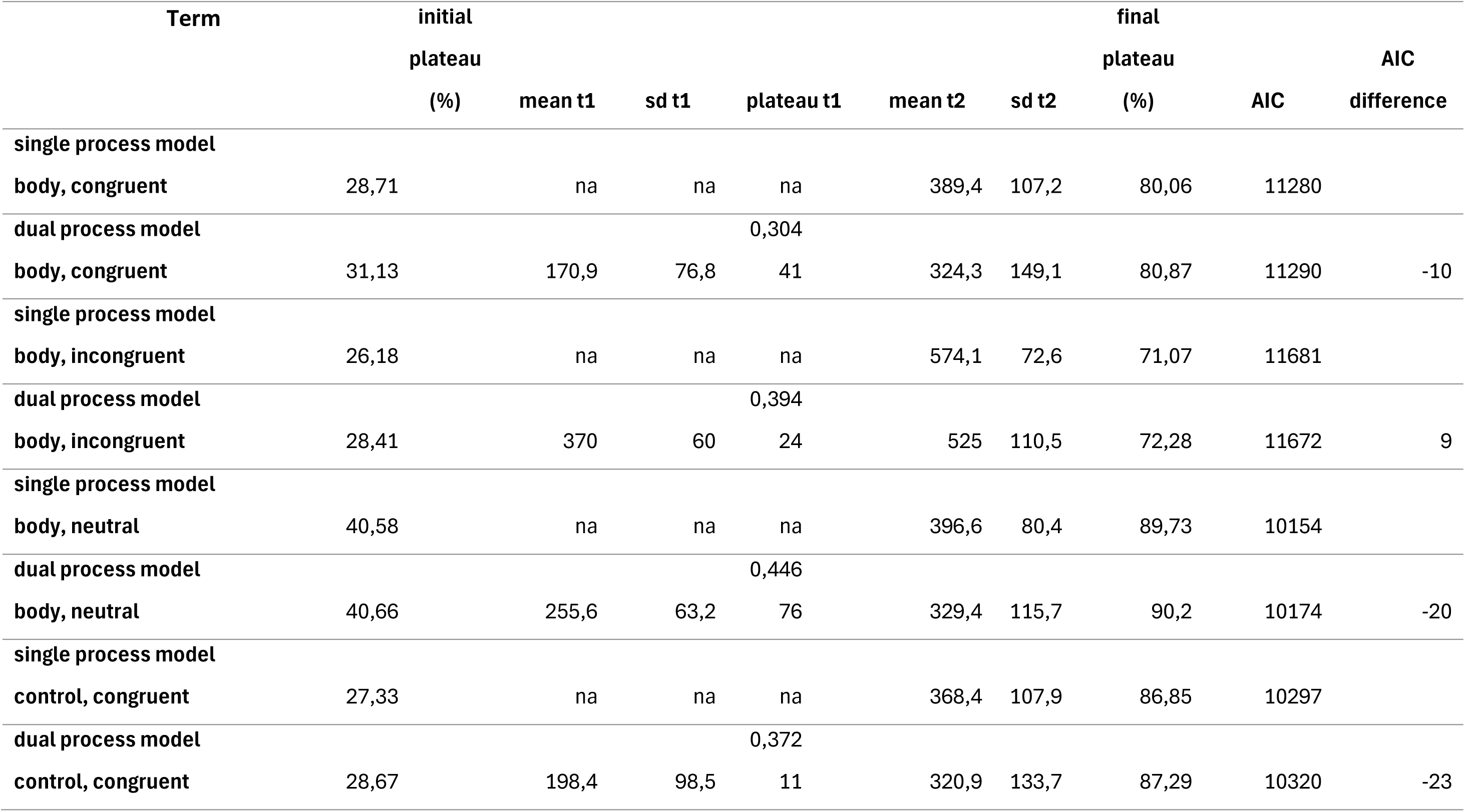
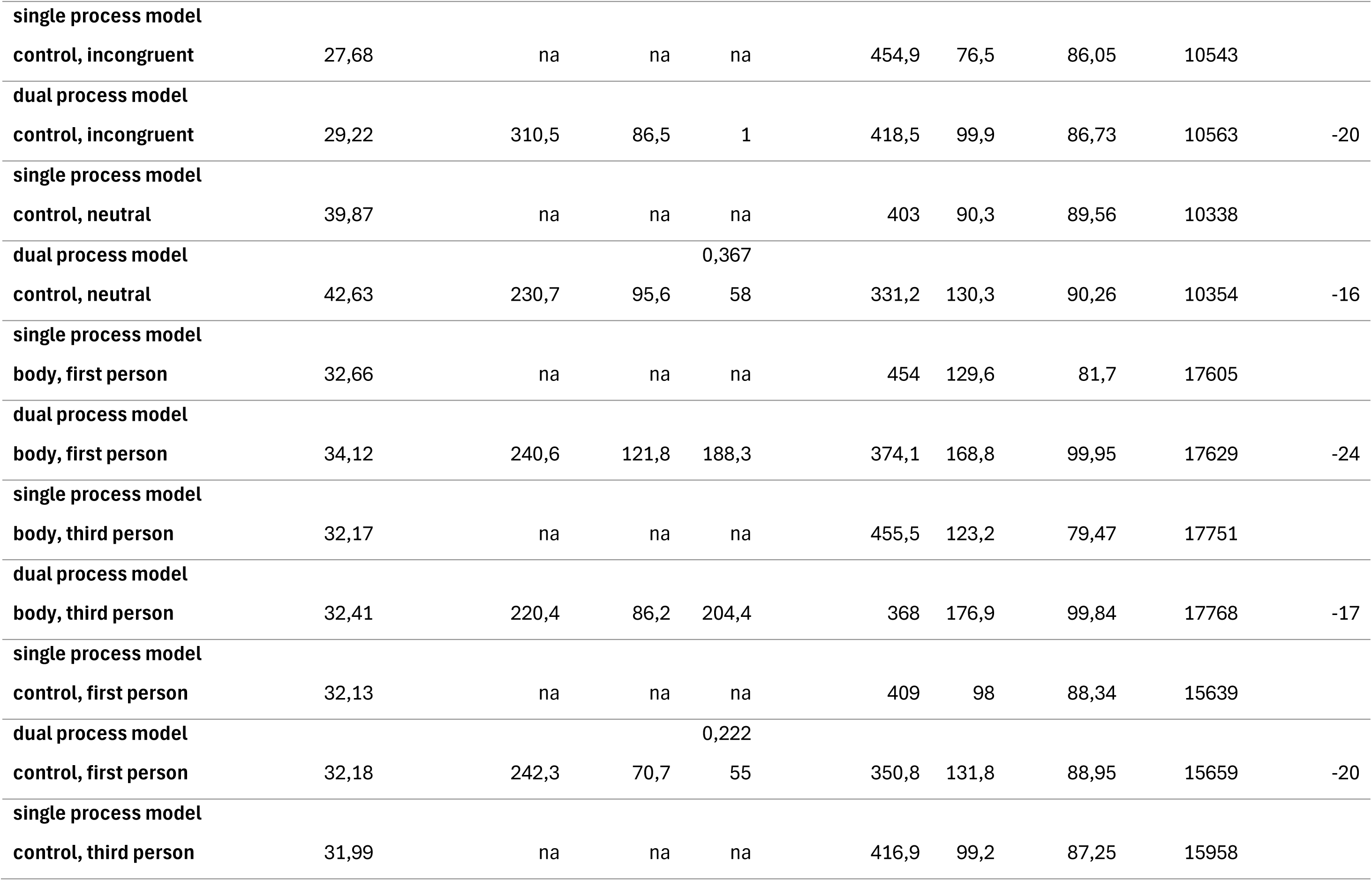
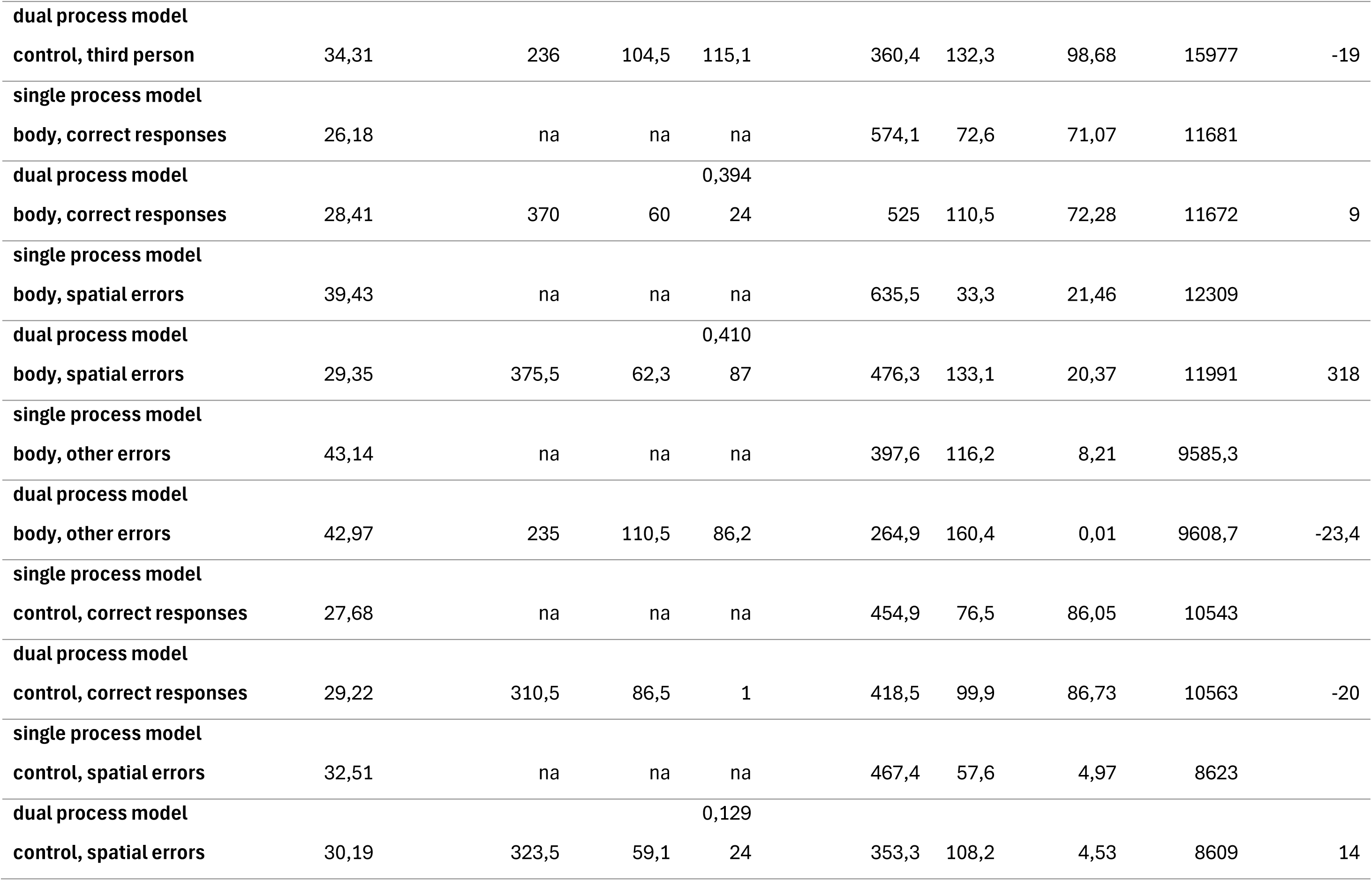
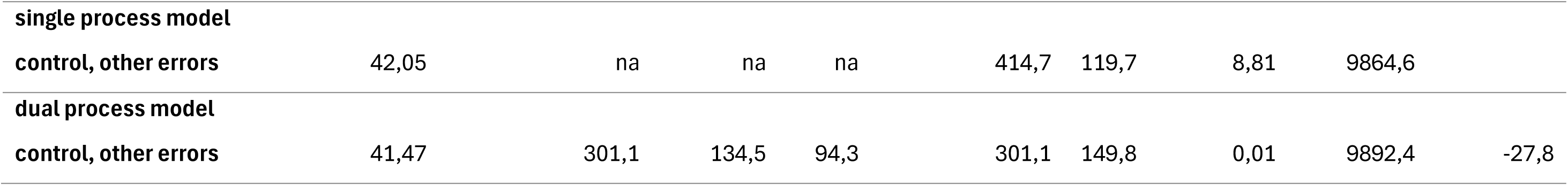
Tables with the values of the various parameters for all the models. Column 1 corresponds to the inflection point for the decrease in the proportion of correct responses and is only present for the complex model. Column 2 corresponds to the variance between participants in column 1. Column 3 corresponds to the inflection point for the increase in the proportion of correct responses. Column 4 corresponds to the variance between participants in column 3. column 5 corresponds to the final plateau calculated by the model. column 6 corresponds to the initial plateau calculated by the model. column 7 corresponds to xxx and is only present for the complex model. Column 8 is the AIC calculated for each model. Column 9 is the difference between the AIC of the simple model and the complex model. A negative value indicates a favour for the simple model, while a positive value indicates a favour for the complex model.

**Table S2.**
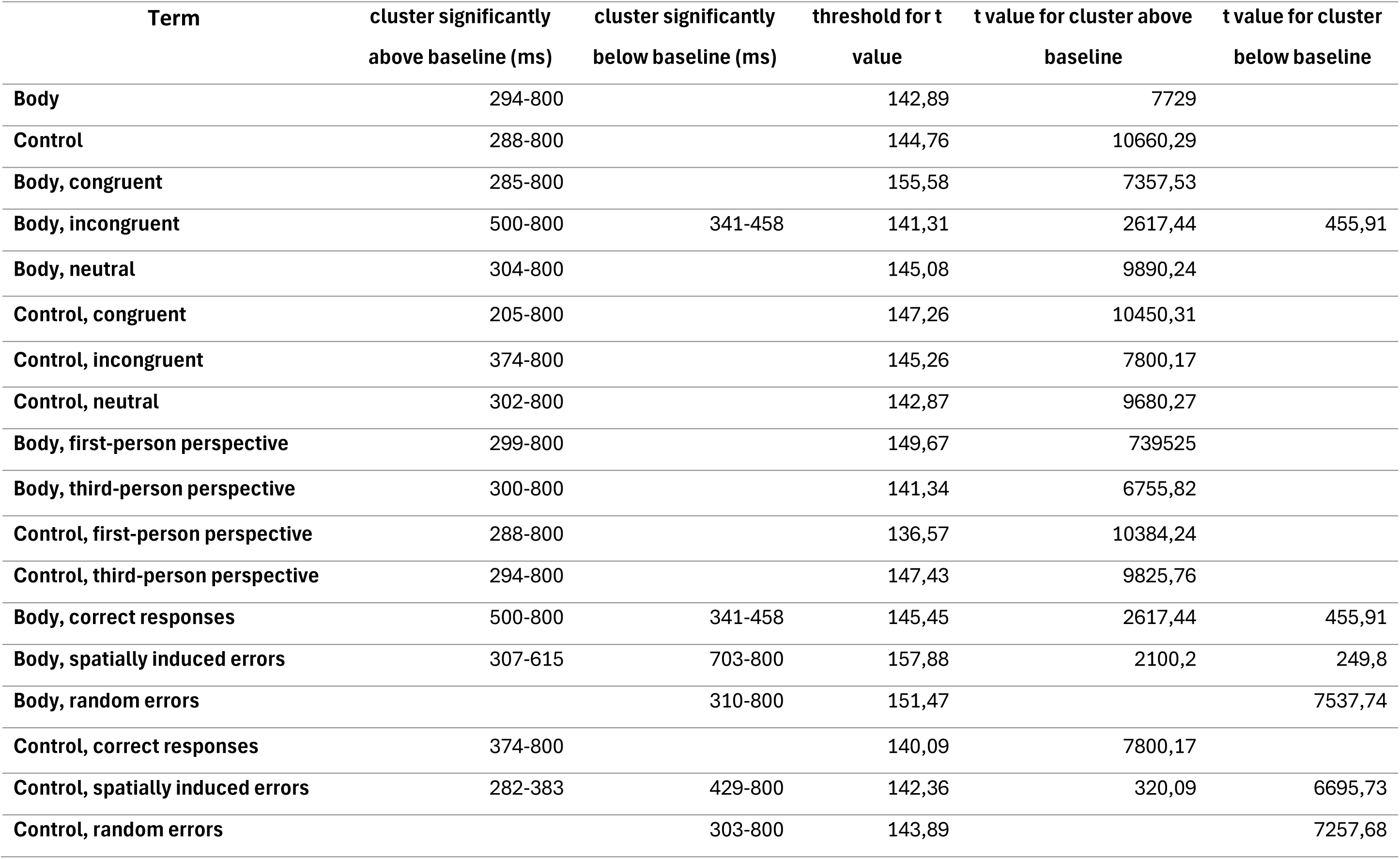
Clusters significantly different from baselines for the different conditions. In column 1 and 2 are the clusters respectively significantly above and below the baseline. Column 3 is the t-value that a cluster must reach to be significant. And columns 4 and 5 are the t-values of the different clusters identified in column 1 and 2 respectively.

**Table S3.**
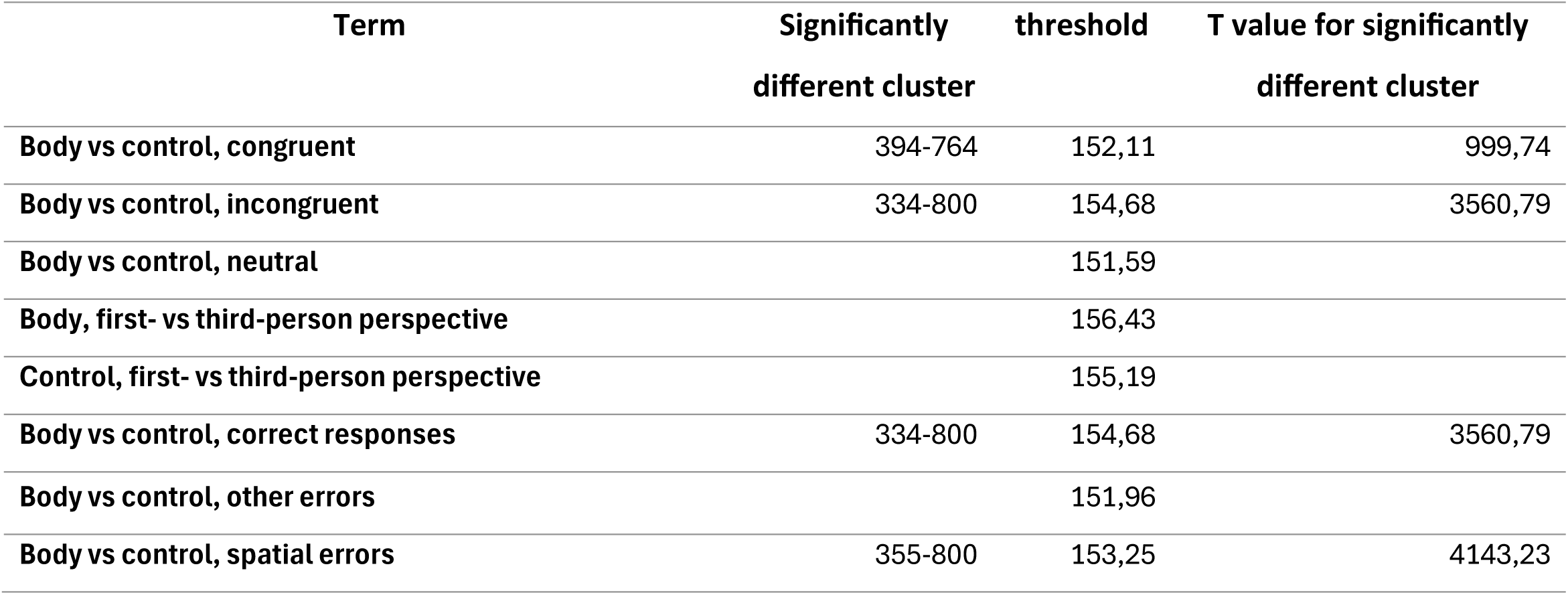
Cluster identified as significantly different by the SMART method for the comparison of two conditions. Column 1 corresponds to the significantly different cluster. Column 2 is the t-value that a cluster must reach to be significant. And column 3 is the t-values of the different clusters identified in column 1

